# Mitochondrial membrane potential regulates nuclear DNA methylation and gene expression through phospholipid remodeling

**DOI:** 10.1101/2024.01.12.575075

**Authors:** Mateus Prates Mori, Oswaldo Lozoya, Ashley M. Brooks, Dagoberto Grenet, Cristina A. Nadalutti, Birgitta Ryback, Kai Ting Huang, Prottoy Hasan, Gyӧrgy Hajnóczky, Janine H. Santos

## Abstract

Maintenance of the mitochondrial inner membrane potential (ΔΨM) is critical for many aspects of mitochondrial function, including mitochondrial protein import and ion homeostasis. While ΔΨM loss and its consequences are well studied, little is known about the effects of increased ΔΨM. In this study, we used cells deleted of *ATPIF1*, a natural inhibitor of the hydrolytic activity of the ATP synthase, as a genetic model of mitochondrial hyperpolarization. Our data show that chronic ΔΨM increase leads to nuclear DNA hypermethylation, regulating transcription of mitochondria, carbohydrate and lipid metabolism genes. Surprisingly, remodeling of phospholipids, but not metabolites or redox changes, mechanistically links the ΔΨM to the epigenome. These changes were also observed upon chemical exposures and reversed by decreasing the ΔΨM, highlighting them as hallmark adaptations to chronic mitochondrial hyperpolarization. Our results reveal the ΔΨM as the upstream signal conveying the mitochondrial status to the epigenome to regulate cellular biology, providing a new framework for how mitochondria can influence health outcomes in the absence of canonical dysfunction.

**Highlights:**

- Mitochondria hyperpolarization leads to nuclear DNA hypermethylation
- DNA methylation regulates expression of mitochondrial and lipid metabolism genes
- Phospholipid remodeling mediates the epigenetic effects of mitochondrial hyperpolarization

## INTRODUCTION

The mitochondrial inner membrane potential (ΔΨM) is primarily maintained by the proton pumping activity of the electron transport chain (ETC) although it can rely on ATP hydrolysis by reverse rotation of the F1 subunit of the ATP synthase under conditions of stress^1^. Maintenance of the ΔΨM is critical for mitochondrial function as mitochondrial protein import, ion homeostasis and many other biochemical processes that occur in these organelles largely depend on it^2–4^. There is a vast body of work describing the consequences of loss of the ΔΨM, which can range from mitochondrial dysfunction (e.g. caused by loss of protein import, ATP production, or Ca^2+^ homeostasis) to a signal for mitochondrial removal through mitophagy. Conversely, little is known about the effects associated with increases in the ΔΨM. In pathology, higher resting ΔΨM was reported in smooth muscle cells isolated from patients with pulmonary hypertension relative to matched healthy controls and in rodent models of the disease^5^; it was also found in glioblastoma^6^. However, the extent to which the increased resting ΔΨM contributes to or is a secondary effect of the pathophysiology of these disorders is unknown.

At the organellar level, studies using isolated mitochondria showed that membrane hyperpolarization promotes reactive oxygen species (ROS) production, which seems threshold-dependent^7^. Higher ΔΨM is known to facilitate mitochondrial Ca^2+^ uptake, activating bioenergetics by its effects on tricarboxylic acid (TCA) cycle dehydrogenases^8^. At the cellular level, deletion of IF1 (or ATPIF1 – *ATP5IF1* official gene symbol), a natural inhibitor of the hydrolase activity of the ATP synthase^9,10^, increased cell viability caused by ETC inhibition and supported cell proliferation during mitochondrial DNA (mtDNA) loss, conditions that depolarized the ΔΨM in wild-type (WT) cells but not in the KO counterparts^11,12^. Most recently, the ΔΨM was shown to impact cell cycle progression, although this seems independent of ROS^13^, and to be modulated by intracellular stimuli. For instance, the growth of cells in phosphate-depleted conditions led to mitochondria hyperpolarization in yeast and in mammalian cells, in an ETC-dependent or independent manner. The higher ΔΨM restored deficient mitochondrial protein import in a mutant yeast strain^14^. This collective body of work indicates that the ΔΨM itself can affect diverse biological outcomes. Also, it seems responsive to intracellular signals and environmental cues, leading to the hypothesis that it can be modulated to influence cellular outcomes. Nonetheless, how cells adapt to chronic rises in ΔΨM and the broader effects associated with it remain largely unknown.

In this study, we used cells genetically depleted of IF1 as a model to study the effects associated with a chronic state of higher resting ΔΨM. Our data highlight the pervasive molecular and genomic changes associated with chronic mitochondrial hyperpolarization, which involves epigenetic remodeling and transcriptional regulation, including of ETC genes. Notably, these effects were driven by the remodeling of phospholipids, reversed by decreasing the ΔΨM and were recapitulated using chemicals to which humans are environmentally exposed, highlighting a new mechanism through which mitochondria may influence health and disease outcomes that has so far been ignored.

## Results

### Chronic loss of IF1 supports a model of increased resting ΔΨM

To define the suitability of using cells depleted of IF1 as a model of chronic mitochondrial hyperpolarization, we started by characterizing the resting ΔΨM of HEK293 IF1-KO vs. WT cells^12^. Using intact cells, TMRE (tetramethylrhodamine ethyl ester) that loads into mitochondria based on the ΔΨM and MitoTracker Green (MTG) to normalize data to mitochondrial content, we found that IF1-KO cells had higher resting ΔΨM than WT controls (Fig. 1A). Similar results were observed when deleting IF1 in the 143B osteosarcoma cell line (Fig. S1A and B). The rise in resting ΔΨM in HEK293 IF1-KO cells was confirmed using permeabilized cells, in which the plasma membrane was removed and mitochondria were energized by adding the complex II substrate succinate. In these experiments, we followed concomitantly the ΔΨM using the fluorescent indicator tetramethylrhodamine methyl ester (TMRM), and the clearance of Ca^2+^ added to the cytoplasmic buffer with FuraFF. The ΔΨM was higher in KO relative to WT cells (Fig 1B upper panel), which was paralleled by faster clearance of cytosolic Ca^2+^ into the mitochondria (Fig. 1B lower panel). Unlike previously reported^15^, we found no changes at the protein levels of the different subunits of the mitochondrial Ca^2+^ uniporter complex (mtCU), which mediate the import of Ca^2+^ in these organelles (Fig. S1C); we also did not find changes in total or phosphorylated AMPK (Fig. S1D). Our data are thus consistent with the higher ΔΨM facilitating the faster uptake of cytosolic Ca^2+^ into the mitochondria of the IF1-KO cells.

**Figure 1.**
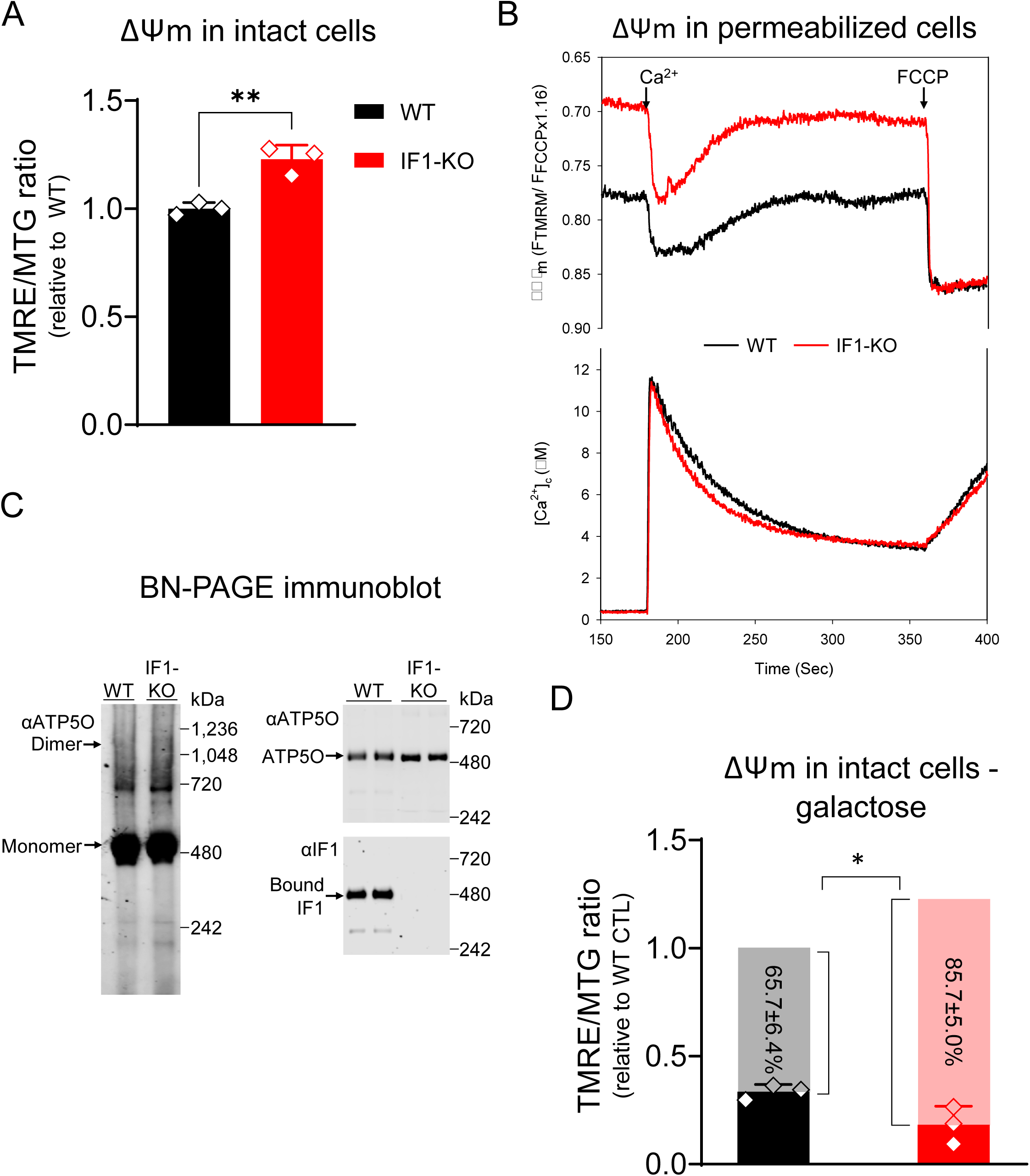
Ablation of IF1 increases resting ΔΨM via ATP synthase hydrolytic activity. A) ΔΨM measured in intact cells using the ΔΨM-sensitive dye TMRE; data were normalized to mitochondrial content using the ΔΨM-insensitive dye MitoTracker® Green (MTG). On *y-*axis, TMRE/MTG ratio, black bar, ΔΨM in WT cells, and red bar, ΔΨM in IF1-KO cells (n = 3, ** *p* < 0.01). Statistical difference by Student’s t test. B) Representative trace of concomitant ΔΨM and Ca^2+^ buffering measurements in permeabilized cells energized with succinate. The black line represents WT cells and the red line IF1-KO cells. FCCP was added at the end of the experiment to fully depolarize the ΔΨM and release the intramitochondrial pool of Ca^2+.^ C) Representative immunoblots of blue native (BN)-PAGE. Left panel, digitonin-permeabilized WT and IF1-KO mitochondria blotted with ATP synthase peripheral stalk subunit OSCP antibody (αATP5O). Complex V monomer and dimer are pointed by the arrows. Right upper panel, WT and IF1-KO mitochondria blotted with αATP5O. Right lower panel, WT and IF1-KO mitochondria blotted with IF1 antibody (αIF1). D) Bar graph of ΔΨM as measured in A in cells grown in galactose 5 mM. *Y* axis depicts the TMRE/MTG ratio, black bar ΔΨM in WT cells, and red bar, ΔΨM in IF1-KO cells. Grey and pink bars represent the ΔΨM difference between WT and IF1-KO grown in high glucose vs. in galactose (n = 3, * *p* < 0.05). Statistical difference by Student’s t test.

While IF1 was thought to bind to the ATP synthase to inhibit its hydrolase activity only under stress conditions, more recent work suggests that heterogenous populations of the enzyme engaged in either ATP synthesis or hydrolysis co-exist in cells^16,17^. In agreement with this, blue native analysis showed that IF1 was also detected in WT cells, but not KO cells, at the same molecular weight of ATP synthase monomers as identified with an antibody against ATP5O (Fig. 1Cl), consistent with previous observations^18^. These data support the notion of IF1 being bound to complex V in WT cells grown under standard cell culture conditions. More importantly, when cells were grown under galactose supplementation that limit the pool of glycolytic ATP that can be hydrolyzed by the ATP synthase, the ΔΨM decreased in WT and KO cells, but the effects were significantly more pronounced in the KO cells (Fig. 1D). Hence, we conclude that the increased resting ΔΨM in the KO cells relies on a significant contribution of ATP hydrolysis under standard cell culture conditions.

### Chronic loss of IF1 leads to a robust transcriptional response

Next, we used RNA-seq to understand the broader cellular adaptations resulting from a higher resting ΔΨM. We found over 6,000 differentially expressed genes (DEGs) between the KO and WT cells, of which 3,884 genes were upregulated (red) and 2,669 were downregulated (blue) (Fig. 2A, see also Table S1). This large transcriptional response in the absence of exogenous stress was unexpected but suggests that the higher resting ΔΨM might provide a signal that cells not only sense but can respond to. Genes associated with glycolysis were upregulated in the IF1-KO cells (Table S1), consistent with unrestrained hydrolysis of glycolytic ATP contributing to the maintenance of a higher resting ΔΨM. The expression of genes involved in transcription, signaling, cell cycle, DNA repair, and lipid metabolism was also increased (Table S1). Interestingly, 8.7% of DEGs in the entire dataset coded for mitochondrial proteins, which were generally downregulated (Table S1). Gene Ontology analysis identified several mitochondrial processes involved in OXPHOS, ETC and the proton motive force as enriched by the downregulated DEGs (Fig. 2B), including 54% of the nuclear subunits of the ETC and 46% of genes involved with the mitoribosome (Fig. 2C). Inhibition of ETC complexes was reported to compensate for oligomycin toxicity as a rheostat mechanism^19^; oligomycin also leads to increased ΔΨM.

**Figure 2.**
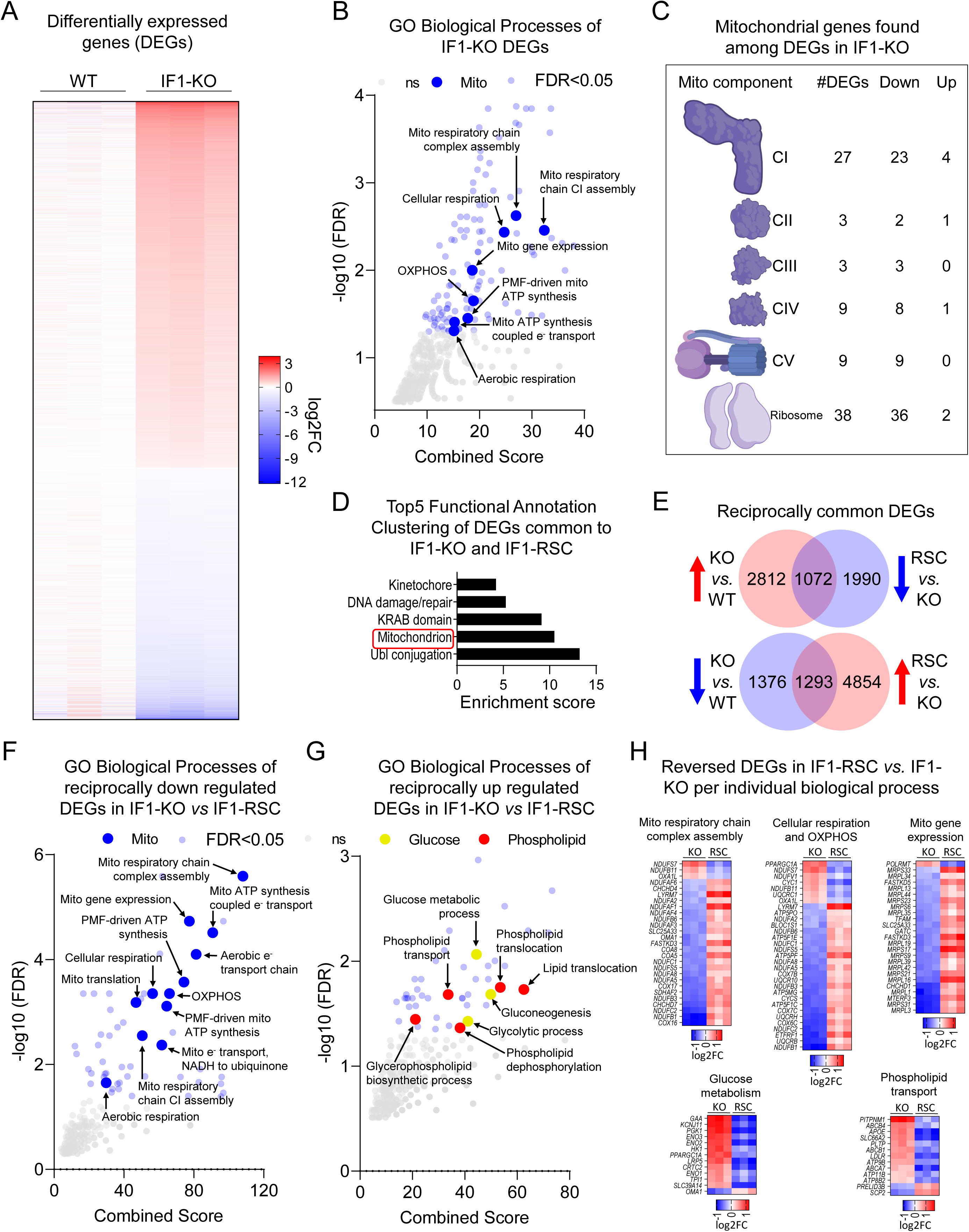
IF1-KO cells with increased ΔΨM engage in a transcriptional feedback loop repressing mitochondrial genetic program. A) Heatmap of differentially expressed genes (DEGs) in IF1-KO vs. WT (False Discovery Rate, FDR < 0.05). The color scheme is presented as a log2 fold change (log2FC). Shades of red represent upregulated genes (log2FC > 0) and shades of blue represent downregulated genes (log2FC < 0). B) Gene Ontology (GO) analysis of DEGs in IF1-KO vs. WT cells. –Log10 FDR values on the y-axis, and combined score on the x-axis. In bigger blue circles, significantly enriched pathways (FDR < 0.05) related to mitochondria. In smaller translucid blue and grey circles, other significantly enriched pathways (FDR < 0.05) and non-significant pathways (FDR > 0.05), respectively. C) Structural depiction of individual complexes of the electron transport chain and mitochondrial ribosomes. The number of DEGs associated with these broad complexes and their transcriptional directionality are also depicted. D) Functional Annotation Clustering of common DEGs reciprocally changed in the inverse direction between IF1-KO vs. WT and IF1-RSC vs. IF1-KO. Keywords and terms found in each cluster are shown on the y-axis and the Enrichment score is on the x-axis. E) Venn diagrams shown the intersection between reciprocally regulated DEGs in IF1-KO vs. WT and IF1-RSC vs IF1-KO. Upper circles: upregulated DEGs based on IF1-KO vs. WT comparison, lower circles represent the downregulated counterparts. F and G) GO analysis of reciprocally regulated DEGs in IF1-KO cells, as depicted and described in B. H) Heatmap of representative pathways reversed in IF1-RSC vs. IF1-KO as identified by GO analysis. Color scheme as described in A. KO is IF1-KO vs. WT, and RSC is IF1-RSC vs. IF1-KO.

We then stably re-introduced IF1 in the KO cells, herein called rescue (RSC) (Fig. S2A) and found over 9K DEGs between the KO and RSC (Table S1). Of these, 4,185 were DEGs between KO vs WT and broadly enriched for mitochondria (Fig. 2D). The expression of 2,365 out of the 4,185 common genes was fully reversed by the re-introduction of IF1 (Fig. 2E). Notably, the 1,293 genes that were downregulated in the KO and upregulated in the RSC enriched for mitochondrial processes while the 1,072 upregulated genes in the KO that were reciprocally downregulated in the rescue enriched for glucose, phospholipid transport and metabolism (Fig. 2F-H). The modulation of phospholipid genes was interesting considering a recent study that linked phosphatidylethanolamine (PE), a main phospholipid of mitochondrial membranes, with regulation of proton conductance by the uncoupling protein 1 (UCP1), at least in brown adipose tissue^20^. Uncoupling through proton dissipation is a known mechanism to regulate the ΔΨM.

The downregulation of mitochondrial genes in the KO cells, which was reversed in the isogenic RSC, prompted us to examine mitochondrial function in the cells using the Seahorse Flux Analyzer. We followed oxygen consumption in glucose– or galactose-supplemented media and found that despite minor changes observed in glucose conditions, no effects were observed when cells were grown under galactose together pointing to no significant mitochondrial respiration impairments (Fig. S2B-C). Likewise, transmission electron microscopy failed to identify changes in mitochondrial ultrastructure (Fig. S2D-E). Thus, we interpret the transcriptional changes associated with the downregulation of genes involved in mitochondrial processes, including those of the ETC, as adaptations to maintain the ΔΨM at an ‘optimal’ state that supports proper mitochondrial and cellular function. In agreement with this notion, we found that the mitochondria of KO cells could further hyperpolarize when they were put under phosphate depleted conditions (Fig. S2F), recapitulating the effects recently reported for other cells^14^. Thus, at baseline, the ΔΨM of KO cells is not maintained at its maximal polarized state.

### Chronic rise in ΔΨM leads to nuclear DNA hypermethylation regulating gene expression

It is increasingly evident that mitochondria communicate with the epigenome^21^, including by modulating DNA methylation that in turn regulates transcription^22,23^. To define whether the broad transcriptional responses found in the KO cells were associated with changes in DNA methylation, we profiled the nuclear genome methylation status at a single nucleotide resolution. Remarkably, we found that the nuclear DNA of the KO cells was significantly hypermethylated relative to the WT counterparts, with 28,832 differentially methylated loci (DML) between them (Table S2). We then focused on promoter methylation to start establishing relationships between this epigenetic mark and transcription. In a simplistic model, promoter hypermethylation is associated with gene repression while promoter hypomethylation leads to the upregulation of genes. We identified promoters based on genomic coordinates of the transcription start site (TSS) +/− 1,500 bases (Fig. S3A) and found 9,420 gene promoters hypermethylated and 3,919 hypomethylated in the KO cells (Table S2). Overlaying DML and DEG lists revealed that 3,629 were differentially methylated and expressed genes (DMEGs, Table S3), out of which 933 had the promoter hypermethylated and were repressed (Fig. 3A, cluster 2) while, 626 genes had the promoter hypomethylated and were upregulated (Fig. 3A, cluster 3). Most notably, genes from cluster 2 collectively enriched for mitochondrial processes (Fig. 3B) whereas those from cluster 3 enriched for phospholipid transport and carbohydrate phosphorylation (Fig. 3C). Collectively, these data suggest that the differential nuclear DNA methylation regulated the transcriptional response to the increased resting ΔΨM in IF1-KO cells.

**Figure 3.**
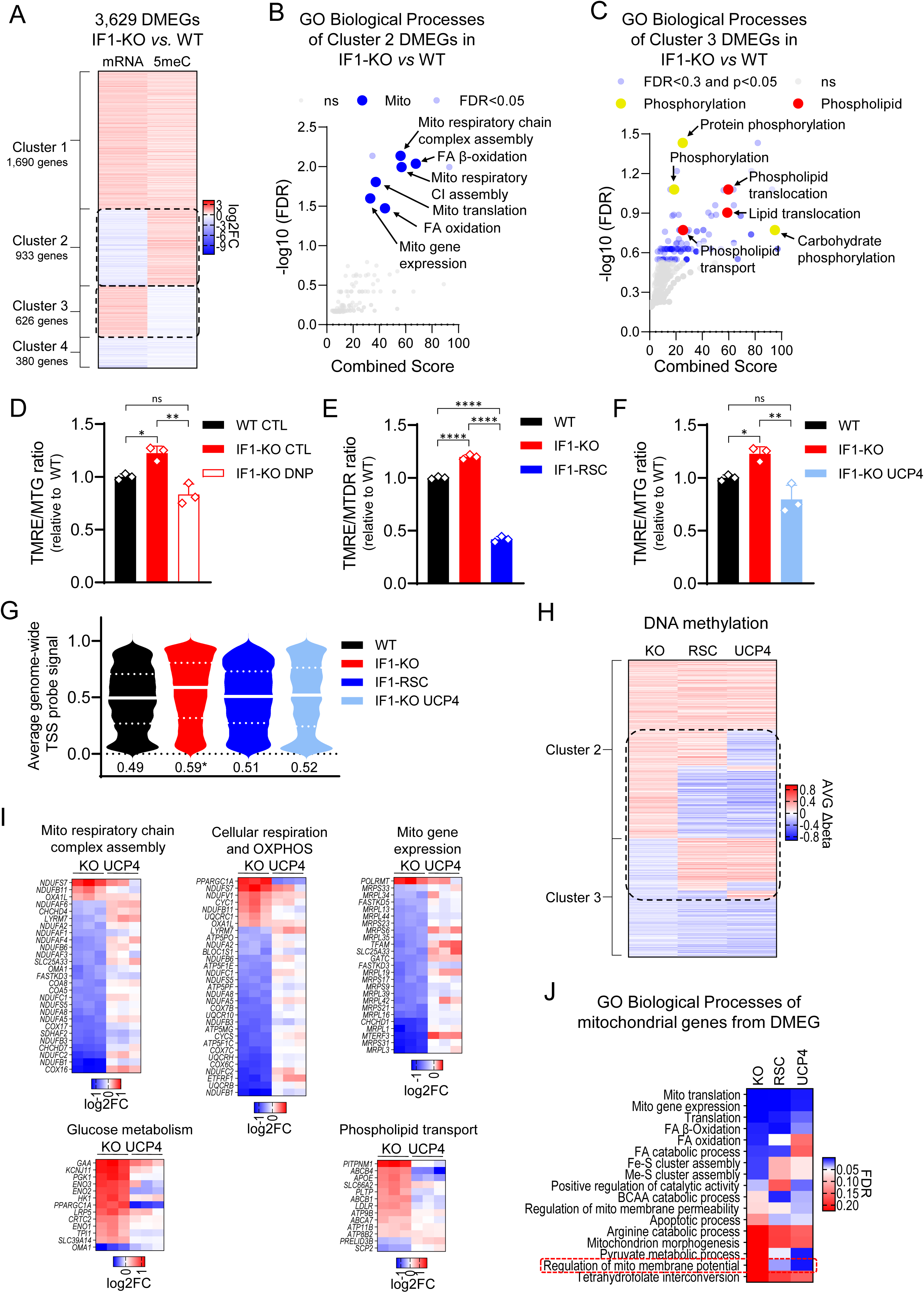
Nuclear DNA is hypermethylated in IF1-KO cells regulating a mitochondrial gene expression program. A) Heatmap of commonly differentially methylated and expressed genes (DMEGs). The color scheme is presented as a log2 fold change (log2FC). Shades of red represent upregulated and hypermethylated genes (log2FC > 0) and shades of blue represent downregulated and hypomethylated genes (log2FC < 0). Left track (mRNA) is gene expression from RNA-seq and right track (5meC) is IF1-KO UCP4 vs. IF1-KO. DMEGs were grouped in 4 clusters based on promoter methylation and gene expression change, respectively: 1) hypermethylated and upregulated, 2) hypermethylated and downregulated, 3) hypomethylated and upregulated, and 4) hypomethylated and downregulated. B and C) GO analysis of DMEG of clusters 2 and 3, respectively. –Log10 FDR values on the y-axis, and combined score on the x-axis. D-F) Bar graph of ΔΨM as measured by ΔΨM-dependent dye TMRE normalized to the ΔΨM-independent dye MTG. Y axis depicts the TMRE/MTG ratio, black bar ΔΨM in WT cells, and red bar, ΔΨM in IF1-KO cells. D) Red-bordered bar, ΔΨM in IF1-KO cells treated for 3 days with DNP 25 μM. E) Blue bar, ΔΨM in IF1-RSC cells. F) Light blue bar, ΔΨM in IF1-KO UCP4 expressing cells. Statistical difference by One-way ANOVA with Tukey’s post-test. G) Violin plot showing DNA methylation status across the entire genome in all 4 genotypes; the Y-axis depicts probe values; white lines indicate the median and dotted lines the quartiles. Numerical values below the violin plot are the median for each genotype. H) Heatmap of DMEGs in clusters 2 and 3 from A. The color scheme is presented as average delta beta (AVGΔbeta, see methods). Shades of red represent hypermethylated genes (AVGΔbeta > 0) and shades of blue represent hypomethylated genes (AVGΔbeta < 0). The left track (KO) is IF1-KO vs. WT, the middle track (RSC) is IF1-RSC vs. IF1-KO, and the right track (UCP4) is IF1-KO UCP4 vs. IF1-KO. I) Heatmap of representative pathways as in Fig. 2H. KO is IF1-KO vs. WT, and UCP4 is IF1-KO UCP4 vs. IF1-KO. J) Heatmap of biological processes (as per Gene Ontology analysis) of mitochondrial genes from DMEG of each paired comparison. The color scheme is presented False Discovery Rate (FDR). Left track (KO) is IF1-KO vs. WT, middle track (RSC) is IF1-RSC vs. IF1-KO, and right track (UCP4) is IF1-KO UCP4 vs. IF1-KO.

### Reversal of hyperpolarization reinstates the epigenetic and transcription landscapes to WT levels

If the higher ΔΨM drives nuclear DNA hypermethylation to regulate transcription, then decreasing the ΔΨM of IF1-KO cells should reverse the epigenetic and transcriptional changes. We pharmacologically decreased the ΔΨM of IF1-KO cells by treatment with the mild protonophore dinitrophenol (DNP, Fig. 3D). We also stably decreased the ΔΨM of IF1-KO cells genetically by re-introducing IF1 (Fig. 3E) and by ectopically expressing UCP4 (*SLC25A27,* Fig. 3F, Fig. S3B and S3C), providing two distinct means to alter the ΔΨM in the cells. UCP4 had been previously shown to mildly decrease the ΔΨM of cells in culture by uncoupling^24^. We then profiled nuclear DNA methylation at a single nucleotide resolution; we focused on the genetic models to avoid potential confounding effects and prevent toxicity associated with long-term exposure to DNP. Strikingly, this analysis showed that KO cells with a lower ΔΨM either through IF1 reintroduction or UCP4 ectopic expression had genome-wide levels of DNA methylation that were close to those observed in the WT counterparts (Fig. 3G, Fig. S3D). Most importantly, these changes were observed at specific loci, including those involved in mitochondrial and phospholipid genes (Fig. 3H, Tables S2 and S3). The ectopic expression of UCP4 also normalized or reversed the expression of specific mitochondrial, glucose metabolism, and phospholipid transport genes (Fig. 3I) essentially mirroring the effects observed upon the re-introduction of IF1 (Fig. 2H). Curiously, DML-containing nuclear-encoded mitochondrial genes enriched for regulation of the ΔΨM, irrespective of the means utilized to decrease it in the KO cells (Fig. 3J). Collectively, these data strongly support the hypothesis that the increased ΔΨM is the upstream signal regulating the epigenetic landscape and transcriptional output of the KO cells.

### ΔΨM-regulated DNA methylation is independent of metabolites or redox changes

Previous work connected dysfunctional mitochondria to nuclear DNA methylation, effects that were primarily driven by modulation of TCA cycle-associated metabolites^21^. Because no changes in mitochondrial function were observed in the KO cells, we questioned whether metabolites were involved with the epigenetic changes observed herein. DNA methylation is kept by opposing reactions that deposit or remove methyl groups from cytosines (Fig. S4A). DNA methyltransferases (DNMTs) use S-adenosylmethionine (SAM) to methylate the DNA, while the removal of that methyl group is initiated by its oxidation by the Ten Eleven Translocation (TETs) enzymes followed by downstream thymine DNA glycosylase-initiated base excision repair^25,26^. TETs are α-ketoglutarate (α-KG)-, oxygen– and iron-dependent dioxygenases^27^ that use ascorbate as a co-factor and are inhibited by their product succinate or the structure-related metabolites fumarate and 2-hydroxyglutarate (2-HG)^28,29^. TETs are also inhibited by oxidative stress^30^ Steady state metabolomics did not reveal significant changes in levels of metabolites that might influence DNMTs activity, including SAM or its related substrate 5′-deoxy-5′-methylthioadenosine (5′-MTA). Similarly, no changes in levels of metabolites associated with the removal of 5-meC by TETs such as α-KG or the competitive inhibitors succinate and fumarate were identified (Fig. 4A). Thus, changes in metabolites that have been associated with DNA methylation and/or the TCA cycle are unlikely to be involved in the epigenetics remodeling identified by modulation of the ΔΨM.

**Figure 4.**
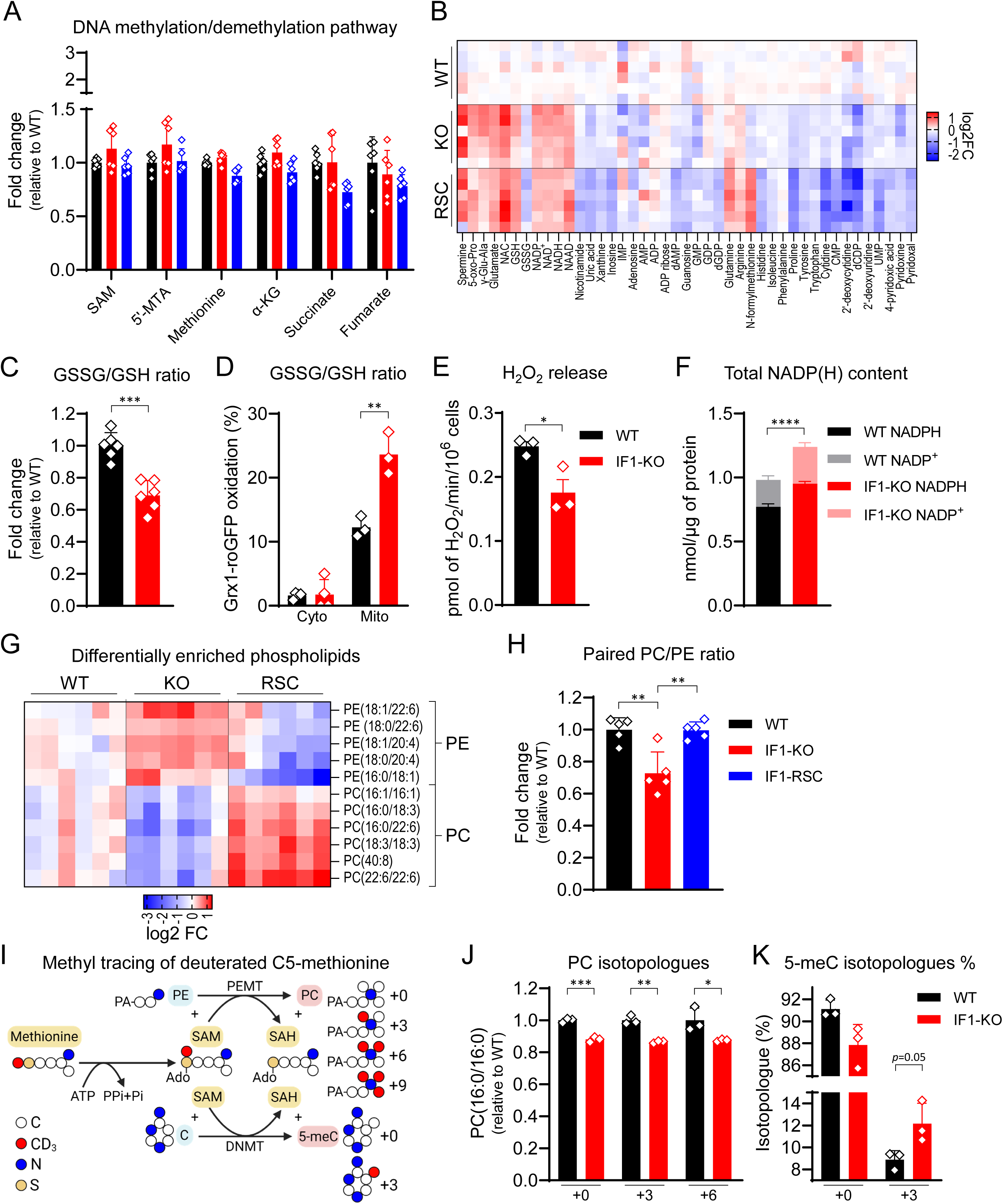
Heads of phospholipids (PC/PE ratio), but not canonical metabolites involved in DNA (de)methylation reactions, better correlate with DNA hypermethylation. A) Measurement of metabolites involved in DNA (de)methylation reactions. Mean WT was set as 1 (n = 6). On the y-axis, fold changes (FC) of metabolites relative to WT. On the x-axis, metabolites. B) Heatmap of differentially enriched metabolites (FDR > 0.05) from steady-state untargeted metabolomics grouped by metabolic pathway. Shades of red represent increased metabolite levels (log2FC > 0) and shades of blue decreased metabolite levels (log2FC < 0). C) Oxidized/reduced glutathione (GSSG/GSH) ratio. On the y-axis, GSSG/GSH ratio relative to WT, n = 6, *** p < 0.001. Statistical difference by Student’s t-test. D) GSSG/GSH accessed by Grx1-roGFP probe ratio in the cytosol (cyto, untargeted) and mitochondria (mito, MTS-containing). On the y-axis, GSSG/GSH ratio relative to WT; n = 3, except IF1-KO, n = 4, ** p < 0.01. Statistical difference by Student’s t-test. E) H_2_O_2_ release accessed by Amplex® Red. On the y-axis, pmol of H_2_O_2_/min/10^6^ cells; n = 3, * p < 0.05. Statistical difference by Student’s t-test. F) Total NADP(H) content. On the y-axis, nmol NADPH or NADP^+^ per μg protein, black and grey bars, NADPH and NADP^+^ in WT cells, respectively, red and pink bars, NADPH and NADP^+^ in IF1-KO cells, respectively (n = 3, ** p < 0.01). Statistical difference by Student’s t-test. G) Heatmap of differentially enriched phospholipids (FDR > 0.05) grouped by head groups. Shades of red represent increased metabolite levels (log2FC > 0) and shades of blue decreased metabolite levels (log2FC < 0). H) PC/PE ratio of paired phospholipids. Four PC/PE pairs (16:0/18:2, 16:0/22:6, 18:0/22:6, and 18:2/18:2) found in metabolomics data were pooled and ratios determined for each sample. Mean WT was set as 1. On the y-axis, fold changes (FC) of metabolites relative to WT; n = 5, ** p < 0.01. Statistical difference by One-way ANOVA with Tukey’s post-test. I) Schematic diagram for tracing methyl group using deuterated C5-methionine. Carbon (C) is depicted in white circles, nitrogen (N) in blue, sulfur (S) in yellow, and deuterated methyl group (CD_3_) in red. Condensation of deuterated C5-methionine with ATP forms deuterated SAM at the methyl-donating position (CD_3_-SAM). Up to three methyl groups of CD_3_-SAM are transferred to PE to form PC by PEMT giving rise to +3, +6, or +9 isotopologues. One methyl group of CD_3_-SAM is transferred to C to form 5meC giving rise to +3 isotopologue. J) Measurement of PC(16:0/16:0) isotopologues. Mean WT was set as 1. On the y-axis, fold changes (FC) of PC(16:0/16:0) relative to WT. On the x-axis, isotopologues (+0, +3, and +6). Black bars FC of PC(16:0/16:0) in WT, red bar, in IF1-KO (n = 3, * p < 0.05, ** p < 0.01, and *** p < 0.001). Statistical difference by Student’s t-test for each isotopologue. K) Percentage (%) of 5meC isotopologues. Total 5meC was set as 100% for each sample and isotopologues +0 and +3 were calculated as % from total. On the *y*-axis, relative abundance of each isotopologue. On the x-axis, isotopologues (+0 and +3). Black bars FC of 5meC in WT, red bar, in IF1-KO (n = 3, p = 0.0505). Statistical difference by Student’s t-test for each isotopologue.

It is conceivable that TET function may be inhibited in the KO cells as a rise in ΔΨM is expected to increase ROS production. However, several lines of evidence indicate that IF1-KO cells have adapted to the chronic state of higher resting ΔΨM by reducing, instead of oxidizing, the cellular environment. For example, levels of glutathione, NADH, NAD^+^, and NADP were increased in the KO cells (Fig. 4B-D), with the ratio of oxidized (GSSG) to reduced (GSH) glutathione decreased (Fig. 4C). Targeting the redox probe GRX1-roGFP2 to the cytosol or mitochondria of WT and KO cells revealed that the mitochondrial matrix, but not the cytosol, was oxidized (Fig. 4D), collectively being inconsistent with the cellular compartments directly continuous with the nucleus being under a state of oxidative stress. Also, levels of the redox couple NADP^+^/NADPH were increased (Fig. 4F) and the amount of hydrogen peroxide (H_2_O_2_) released in the medium as gauged by Amplex® Red was decreased in the KO cells (Fig. 4E). Altogether, these data point to IF1-KO cells adapting to the chronic rise in resting ΔΨM by increasing their antioxidant capacity.

Further inspection of the metabolomics data identified that phospholipids were not only modulated in the KO cells but their levels were fully reversed in the RSC (Fig. 4G). Specifically, levels of PE and phosphatidylcholine (PC), the two major phospholipid components of mitochondrial membranes, were altered such that they affected the global cellular PC/PE ratio (Fig. 4H). From an epigenetic perspective, a decrease in the cellular PC/PE ratio caused by the deletion of phosphatidylethanolamine methyltransferase (PEMT), the enzyme that methylates PE into PC in one of the pathways of PC production, was shown to cause histone hypermethylation in yeast^31^. Nevertheless, under our experimental conditions, histones were not hypermethylated in any of the genotypes analyzed (Fig. S4B), suggesting that in mammalian cells methylation of the DNA, not histones, is primarily affected by a decrease in the cellular PC/PE ratio. We tested this hypothesis by following the fate of methyl groups using L-methionine-(*methyl*-d3), a methionine isotopologue deuterated at C5 from which carbons will eventually be incorporated into PC and the DNA (Fig. 4I). If methylation of PE was reduced in the KO cells, we expected decreased levels of +3– or +6-labeled PC along with increased +3 labeling of cytosine. Remarkably, concomitant to the decreased labeling of PC (Fig. 4J) were increases in labeling of 5-meC (Fig. 4K), strongly supporting the hypothesis that the decreased PC levels and increased DNA methylation in the IF1-KO cells derive from less PE methylation. It is worth noting that we were able to detect +3 and +6 isotopologues of PC(16:0/16:0), but not +9. This is consistent with methionine been recycled via homocysteine with methyl donor 5-methyltetrahydrofolate through serine-glycine one-carbon metabolism^32^. Indeed, low levels of +0 methionine were detected in the WT and IF1-KO (Fig. S4C), even though the only exogenous source of methionine was the deuterated (+3) form.

### Changes in phospholipids underlie the effects of nuclear DNA methylation by the ΔΨM

If changes in PC and PE levels are an adaptation to higher resting ΔΨM that mediate DNA hypermethylation, then increasing the ΔΨM of WT cells should also change the PC/PE ratio. To test this, we first identified chemicals that can chronically increase the ΔΨM without changes in mitochondrial volume or toxicity. To this end, we took advantage of data provided by Tox21, a consortium of different US Federal Agencies that aims at developing better toxicity assessment methods to evaluate the safety of chemicals, pesticides, food additives, contaminants, and medical products. In one phase, Tox21 leveraged high-throughput screenings, using various cell types and assays, to evaluate the activities of 10K chemicals. ΔΨM was among the parameters evaluated using the ratiometric fluorescent dye JC-10. It was found that about 30% of the 10K chemicals screened changed JC-10 fluorescence ratio, out of which ∼5% increased the ΔΨM^33^. We selected a few of the chemicals, including pharmaceutical drugs, to first confirm their ability to increase the ΔΨM using TMRE, normalizing data to mitochondrial content based on MitoTracker^TM^ Green. The chemicals/drugs were selected because of the potential for long-term human exposures, which would provide a physiological setting in which the resting ΔΨM may be chronically increased through voluntary, occupational or contaminant environmental exposures. In all cases, the test articles caused acute mitochondrial hyperpolarization (Fig. 5A and Fig. S5).

**Figure 5.**
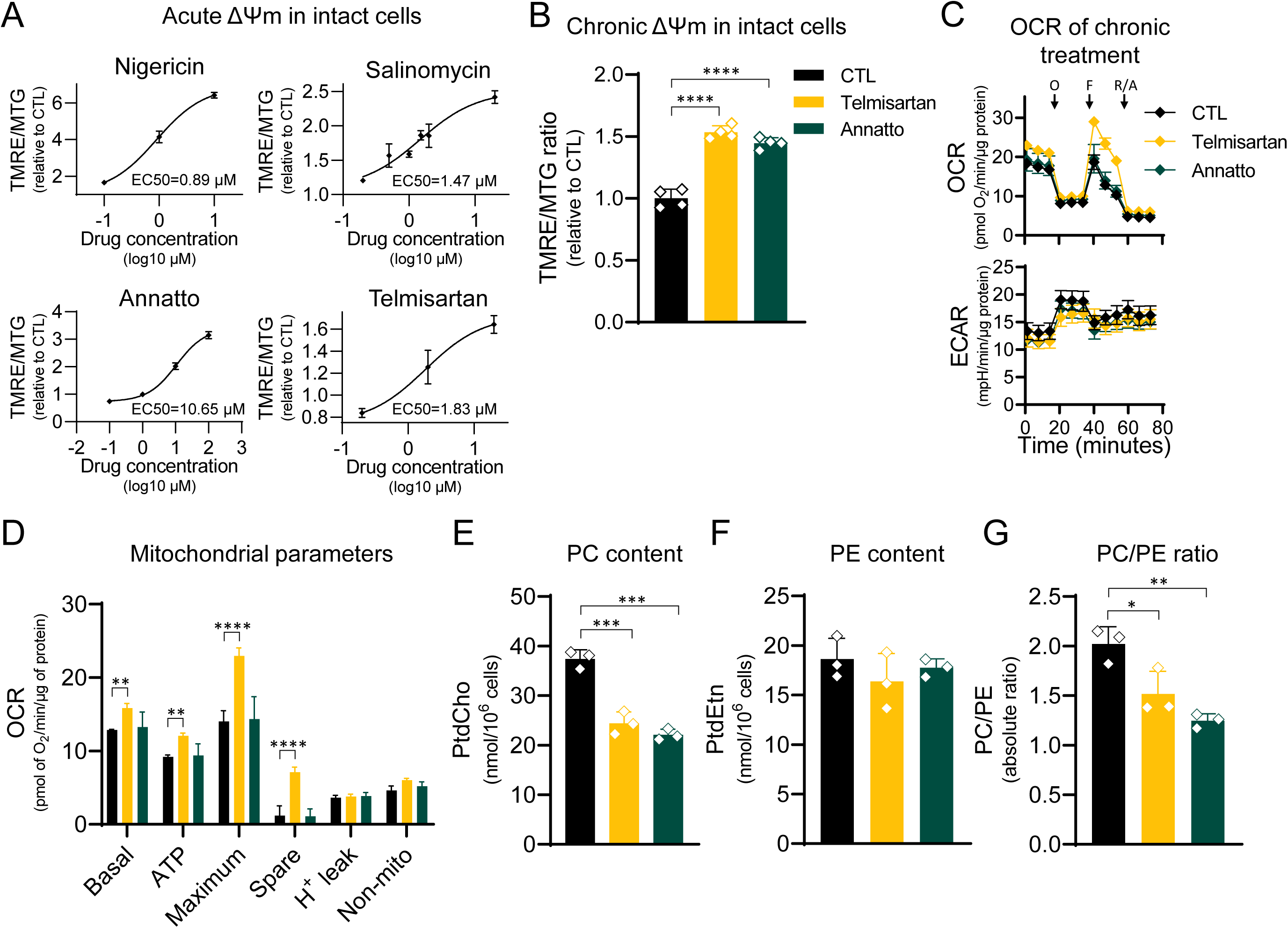
Chronic drug-induced ΔΨM hyperpolarization recapitulates IF1-KO PC/PE shift. A) Dose-response Hill slope was used to determine EC50 of drugs tested. On the y-axis, the TMRE/MTG ratio, and on the x-axis drug concentration in log10 μM. B) Drug-induced ΔΨM hyperpolarization in cells treated for 10 days; n = 4, **** p < 0.0001. C) Upper graph, oxygen consumption rate (OCR) on the y-axis in pmol O_2_/min/μg protein. Lower graph, extracellular acidification rate (ECAR) on the y-axis in mpH/min/ μg protein. Arrows indicate injection of mitochondrial inhibitors: oligomycin A 1 μM (O), FCCP 1 μM (F), and rotenone and antimycin A 1 μM (R/A) (n = 3). D) Mitochondrial parameters: Basal mitochondrial respiration, ATP-linked respiration, maximum respiration, spare respiratory capacity, H^+^ leak, and non-mitochondrial respiration; n = 3, ** p < 0.01, **** p < 0.0001. E and F) PC and PE content in cells chronically treated with ΔΨM hyperpolarizing drugs; n = 3, *** p < 0.01. G) Absolute PC/PE ratio in cells chronically treated with ΔΨM hyperpolarizing drugs; n = 3, *** p < 0.01. In all panels, statistical difference was tested by one-way ANOVA with Dunnett’s post-test.

Next, we performed time-course experiments to establish doses to which WT cells could be chronically exposed to these chemicals without toxicity, thus allowing them to adapt to the rise in resting ΔΨM. Telmisartan, an anti-hypertensive drug, and annatto, a food additive, proved to be well-tolerated for 10 consecutive days without signs of toxicity (i.e., loss of cell viability). On day 10, we analyzed the ΔΨM, PC, and PE content of WT cells treated with vehicle or with telmisartan or annatto. Exposure to both drugs chronically increased the resting ΔΨM of WT cells (Fig. 5B). Similar to the IF1-KO cells, the rise in resting ΔΨM was not associated with mitochondrial function as judged by oxygen consumption and extracellular acidification (Fig. 5C). More importantly, analysis of PC/PE content showed that the cellular ratio also decreased, paralleling both the ΔΨM increases (Fig. 5D) and the effects observed in IF1-KO cells. Thus, modulation of phospholipids seems to be a hallmark adaptation to chronic increases in the ΔΨM. Together with the lack of mitochondrial dysfunction identified in the IF1-KO cells, these data demonstrate a novel means through which mitochondria may significantly influence health outcomes without the presence of dysfunction.

## Discussion

In this paper, we established that modulation of the ΔΨM is mechanistically linked to nuclear DNA methylation. We provide several lines of evidence to support that phospholipid remodeling, and not metabolites, underlies the epigenetic changes. Our data also reveals a large-scale adaptation in reprogramming of transcription, including of ETC and phospholipid metabolism genes, as resulting from chronic mitochondrial hyperpolarization. Remarkably, the loci of these genes are also the ones that are differentially methylated at the DNA level. Even though metabolic rewiring was observed in the IF1-KO cells, it did not involve classic metabolites previously associated with DNA methylation but rather those associated with redox homeostasis, including through the pentose phosphate pathway (PPP). These are likely adaptations to increases in mitochondrial ROS production due to a higher resting ΔΨM. Activation of the PPP helps explain the previously reported effects of the ΔΨM on cell proliferation^12,13^.

In both genetic and pharmacological models of ΔΨM hyperpolarization, we found a change in the PC/PE ratio derived primarily from a decrease in PC. Changes in cellular phospholipids through the loss of PE to PC methylation have been previously shown to cause histone hypermethylation in yeast devoid of PEMT or under starvation, which was proposed as a means to recycle SAM to maintain the cellular methionine/methyl cycle^31,34^. In our cells, neither were histones hypermethylated nor SAM accumulated, suggesting fundamental differences in the epigenetic outcomes associated with PE/PC remodeling in yeast and mammalian cells. As the existence of enzymatic-driven DNA methylation in yeast remains debatable^35^, it is possible that in mammalian cells the DNA rather than histones is the acceptor of PE-derived methyl groups. Intuitively, the DNA would seem a more efficient way to maintain the recycling of methyl groups from SAM to s-adenosyl-homocysteine (SAH) given the abundance of cytosines (2 billion) over that of histones (∼240 million) in the mammalian genome^36^. The vast source of recycling sites at the DNA level may also explain the lack of accumulation of SAM in the IF1-KO cells. Alternatively, the distinct epigenetic outcomes may reflect differences associated with the stimulus i.e., chronic ΔΨM increases (our study) vs PEMT deletion or cellular starvation^31,34^, or with the supraphysiological accumulation of SAM associated with complete loss of PEMT activity. Irrespective, why PEMT activity, which is regulated by substrate availability or transcriptionally^37^, seems to respond to a rise in resting ΔΨM remains to be determined.

That a chronic rise in the ΔΨM leads to phospholipid remodeling is clear based on our data, but why such remodeling occurs remains to be fully defined. In brown adipose tissue, PE in the mitochondrial membrane was recently shown to regulate proton flux through UCP1, presumably affecting the ΔΨM although this was not measured^20^. While our cells do not express UCP1, we speculate that an increase in the PE content in the mitochondria membrane of IF1-KO cells could be deployed in a similar fashion to allow maintaining an optimal hyperpolarized state supporting proper mitochondrial and/or cellular function. A schematic depiction of our working model is presented in Fig. 6. We propose that increased ΔΨM first triggers mitochondrial PE/PC remodeling to support a sustained hyperpolarized state by the inner mitochondrial membrane. This in turn affects the overall cellular PC/PE ratio, which not only provides the methyl groups to methylate the nuclear DNA and rewire transcription, but becomes the signal that initiates such remodeling.

**Figure 6.**
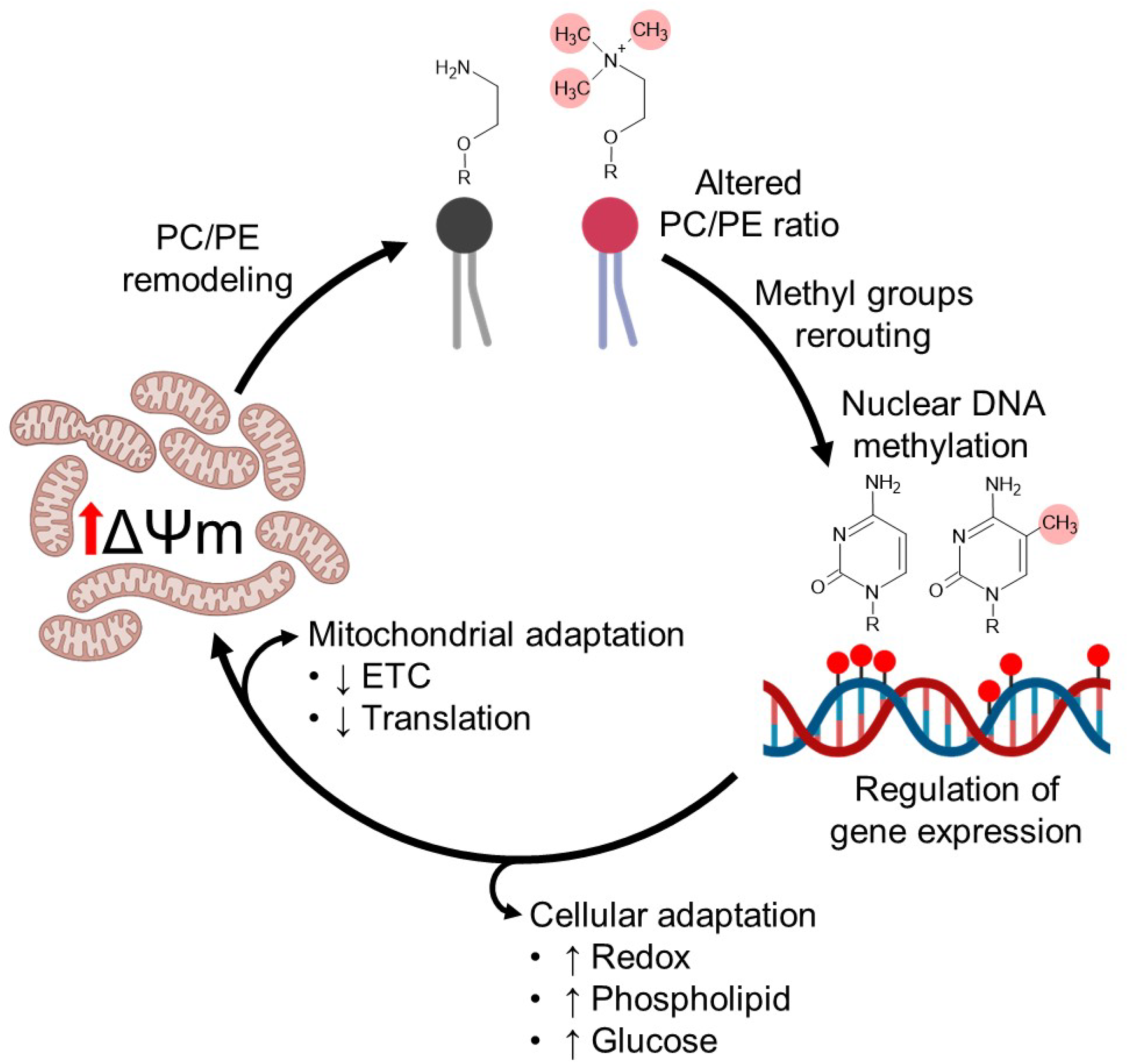
Cellular and mitochondrial adaptations to a chronic state of mitochondrial hyperpolarization. Chronic ΔΨM hyperpolarization leads to mitochondrial membrane phospholipid remodeling with increased PE, which can broadly help regulate proton flux and thus the ΔΨM. This increased membrane PE derives from its decreased methylation into PC, not only impacting PC levels but the PC/PE cellular ratio. Methyl groups from SAM not used to generate PC are then re-routed to the nuclear DNA, including over the promoter regions of thousands of genes. The differentially methylated loci in the nucleus regulate an adaptive gene expression program that involve both inhibition of mitochondrial genes, likely to further help regulate proton flux and the ΔΨM, as well as other broader cellular adaptations involving redox homeostasis and glucose/lipid metabolism.

Our studies did not specifically evaluate mitochondrial membrane phospholipids, but it is noteworthy that a recent study revealed the interplay between cellular and organellar phospholipid composition. It was shown that mitochondrial biogenesis stress leads to whole cell phospholipid remodeling, reflecting changes in membrane composition in the peroxisomes, golgi, endoplasmic reticulum and mitochondria themselves – highlighting the interdependence and connectivity of lipid homeostasis within the cells^38^. How the vast cell-wide changes in phospholipids end up regulating DNA methylation and transcription at specific loci when mitochondria are hyperpolarized remains unknown. Specifically, why modulation of ETC and mitochondrial translation genes were pervasive is unclear, but they might be deployed as additional means, beyond PE/PC remodeling, to further prevent excessive hyperpolarization of the inner mitochondrial membrane. Irrespective, the fact that drugs and chemicals lead to similar rewiring raises questions about how inherent (e.g. pulmonary hypertension) or environmentally induced chronic rises in ΔΨM may impact health and disease outcomes.

In summary, our results provide insights into the molecular and genomics adaptations associated with a chronic rise in resting ΔΨM. Our data adds to the growing body of evidence identifying a relationship between phospholipids and mitochondria while revealing that the ΔΨM can serve as the upstream signal to communicate with the epigenome. Notably, this occurs in the absence of mitochondrial dysfunction. Whether small decreases in the ΔΨM can lead to similar phenotypes is unknown. Overall, our data raise fundamental questions about the potential insidious effects of environmental exposures that affect the ΔΨM, including in their potential to influence disease. It also highlights the possibility of modulating the ΔΨM to modify health outcomes.

## Acknowledgements

We are grateful for the technical assistance of staff members of the Epigenomics, Metabolomics and Molecular Genomics Core Facilities at NIEHS, including Gregory Solomon, Nicole Reeves, Jason Malphurs, Laura Wharey and Dr. Kevin Gerrish. We also thank Dr. William Copeland (Genome Integrity and Structural Biology Laboratory, NIEHS) and Dr. Xiaoling Li (Signal Transduction Laboratory, NIEHS) for critical review of the manuscript. This work was funded by intramural research funds (JHS) and grants R01DK125897-03S1 and R01 GM151536-01 to GH.

## Author Contributions

M.P.M. and J.H.S conceptualized the study, designed the experiments, and wrote the manuscript. M.P.M. did all cell culture experiments, including generating the genetic models, ΔΨM measurements and preparing samples for all ‘omics’ approaches and data analyses. O.A.L. performed some RNA-seq experiments; A.B. analyzed the DNA methylation and RNA-seq data; D.G. was involved in Western blots and nucleic acid extractions; C.N. performed the TEM and part of steady state metabolomics; B.R. performed metabolic tracing and its initial analysis; K.T.H. and P.H. performed experiments in permeabilized cells and with roGFP and analyzed data. G.H. revised the manuscript.

## STAR METHODS

### Cell lines

HEK293T cells carrying a tetracycline (Tet)-on inducible DN-POLG and the ATPIF1 KO isogenic derivatives were provided by Dr. Navdeep Chandel (Northwestern University); these cells were previously described^12^. The pcDNA3.1-hUCP4-NE was purchased from Addgene (plasmid # 102362, deposited by Dr. Shu Leong Ho). IF1-KO cells were used to generate isogenic derivatives ectopically expressing IF1 (IF1-RSC) or UCP4 (IF1-KO UCP4). Stable expression was obtained transfecting the cells using Lipofectamine^TM^ 3000 (Invitrogen^TM^) and selecting multiple clones. IF1 was deleted using CRISPR/Cas9 (GenScript) in the 143B osteosarcoma cell line; cells were expanded after single clone selection in DMEM high glucose (D-glucose 4.5 g/L, 25 mm) supplemented with FBS 10%, and G418 0.8 mg/mL. HEK293 cells were maintained in DMEM high glucose (D-glucose 4.5 g/L, 25 mm) supplemented with FBS 10%, pyruvate 1 mM, uridine 50 μg/mL, and 0.16% penicillin/streptomycin, hygromycin 150 μg/mL and, blasticidin 5 μg/mL at 37°C and 5% CO_2_. 143B WT and 143B IF1-KO cells were maintained in DMEM high glucose (D-glucose 4.5 g/L, 25 mm) supplemented with FBS 10%, and 1% penicillin/streptomycin, at 37°C and 5% CO_2_.

### Mitochondrial membrane potential and chemical treatments in intact cells

ΔΨM was accessed using tetramethylrhodamine ethyl ester (TMRE). 5×10^5^ cells were plates in a 6-well plates (5.6×10^4^ cells/cm^2^), and incubated at 37 °C in a 5% CO_2_ humidified incubator for 24 h. Medium was removed and washed once with FBS-free DMEM. Cells incubated in 2 mL FBS-free DMEM with TMRE 25 nM and MitoTracker® Green (MTG) 50 nM at 37 °C for 15 min. Alternatively, MitoTracker® DeepRed (MTDR) 100 nM was used since HEK293T IF1-RSC cells expressed EGFP. To induce mild mitochondrial uncoupling with 2,4-dinitrophenol (DNP, Sigma-Aldrich®) without toxicity, we titrated several concentrations of DNP. Next, 1.25×10^5^ cells were plated in a 6-well plate and incubated at 37 °C in a 5% CO_2_ humidified incubator with 25 µM for 3 days. For phosphate starvation experiments, recently published protocols using phosphate-free medium were followed^14^. Fluorescence was analyzed in BD LSRFortessa™ Cell Analyzer using FITC channel for MTG and PE-Cy7 channel for TMRE. 10,000 events were analyzed. Mean fluorescence values were converted as ratio of TMRE/MTG.

### Mitochondrial Ca^2+^ uptake and membrane potential measurements in permeabilized cells

ΔΨ_m_ was recorded simultaneously with [Ca^2+^]_c_ using a multi-excitation and dual-emission fluorometer (DeltaRAM; Horiba, NewJersey, USA) as described^39,40^. Cells were harvested and washed with cold Na-Hepes-EGTA buffer containing 120 mM NaCl, 5 mM KCl, 1 mM KH_2_PO_4_, 0.2 mM MgCl_2_, and 20 mM Hepes-NaOH, pH 7.4. In 37°C and under stirring condition the same aliquots of cells (1.8mg) were permeabilized using 30-40 μg/ml Digitonin in 1.5 ml intracellular medium buffer (ICM:120 mM KCl,10 mM NaCl,1 mM KH_2_PO_4_, 20 mM Hepes-Tris, pH 7.2) supplemented with 5 μg/ml protease inhibitors leupeptin, antipain and pepstatin for 5 min. In all the experiments, 2 mM MgATP, 2 mM succinate, and 2 μM thapsigargin (Enzo), TMRM (1.5 μM) and 1.0 μM fura2FF (Kd=4.5 μM, TEFLabs) were present. Uncoupler, FCCP and oligomycin, (5 μM and 5 μg/ml, respectively) was applied at the end of each run to dissipate remaining ΔΨ_m_. TMRM was recorded with 540 nm excitation and 580 nm emission, whereas furaFF at 340nm and 380nm excitation and 500nm emission. Calibration for maximum and minimum fura2FF response was performed by adding 2mM Ca^2+^ and 10M EGTA/TRIS pH8.5, respectively.

### Histone purification

Histones were purified according to Shechter et al^41^. Briefly, approximately 5×10^6^ cells were incubated in hypotonic lysis buffer (Tris-HCl 10 mM, pH 8.0, KCl 1 mM, MgCl_2_ 1.5 mM, and DTT 1 mM) on rotator at 4 °C for 30 min. Nuclei were pelleted at 10,000 *g* at 4 °C for 10 min. Supernatant was discarded and nuclei resuspended in 400 μL of H_2_SO_4_ 0.4 M. Acid extraction was performed on rotator at 4 °C overnight. Samples were centrifuged at 16,000 *g* at 4 °C for 10 min. Supernatant was transferred into new tubes and histones precipitated with 132 μL of trichloroacetic acid 100% (w/v). Samples were incubated on ice for 30 min and centrifuged at 16,000 *g* at 4 °C for 10 min. Supernatant was removed and pellet was washed thrice with ice-cold acetone by centrifugation at 16,000 *g* at 4 °C for 5 min. Histones were air-dried and quantified using BCA.

### Native PAGE and Immunoblotting

Native PAGE was performed according to Jha et al.^42^ and transferred according to Brischigliaro et al^43^. Briefly, 50 μg of mitochondrial fractions were thawed on ice for 30 min. Samples were mixed with 8 μL digitonin 5% (w/v), 5 μL 4X NativePAGE sample buffer and a sufficient volume of deionized to 20 μL (subtracting the volume of the mitochondrial fraction). Samples were incubated on ice for 20 min and centrifuged at 20,000 *g* at 4 °C for 10 min. 15 μL of the supernatants transferred into new tubes. 2 μL of Coomassie G-250 sample additive were added per sample, mixed, and 12 μL of each sample were loaded. Electrophoresis in NativePAGE™ 3 to 12% gels was performed in XCell™ SureLock™ with NativePAGE™ (NP) buffer (anode) and Coomassie Brilliant Blue G-250 (CBBG) 0.02% (w/v) in NP buffer (dark blue cathode) at 150 V for 30 min. After 30 min, dark blue cathode buffer was removed and replaced with CBBG 0.002% (w/v) in NP buffer (light blue cathode). BN-PAGE gels were transferred to methanol-activated PVDF membranes in Dunn carbonate buffer (10 mM NaHCO_3_, 3 mM Na_2_CO_3_). Transfer was performed at constant current of 300 mA at 4 °C for 1 h using XCell™ Blot Module (Invitrogen™). Proteins were fixed/denatured with 8% acetic acid for 5 min, and CBBG was removed washing thrice with methanol. Membranes were blocked using EveryBlot Blocking Buffer (Bio-Rad©) and incubated overnight with antibodies listed in table S. Detection was performed in Odyssey® CLx Imager (LI-COR©).

### SDS-PAGE and immunoblotting

Protein samples were solubilized in cold RIPA Lysis and Extraction Buffer supplemented with Halt™ Protease and Phosphatase Inhibitor Cocktail (Thermo Scientific^TM^) and centrifuged at 20,000 *g* for 10 min. The supernatant was mixed with NuPage LDS sample buffer (Invitrogen^TM^), resolved by Tris-glycine SDS-PAGE, and transferred to PVDF or nitrocellulose membrane. Detection was performed in Odyssey® CLx Imager (LI-COR©). Cell lysates (25µg of proteins) were loaded per well. Primary antibodies and dilutions are as follows: MICU1 (Sigma, HPA034780, 1:500), MICU2 (Abcam, ab101465 1:500 and Bethyl, A300-BL 19212, 1:1000), MCU (Sigma Aldrich, AM Ab91189, 1:1000), EMRE (Bethyl, A300-BL 19208,1:1000). Secondary antibodies and dilutions: goat-anti rabbit IRDye 680RD (LI-COR 925-681817, 1:10,000), and goat-anti mouse 680RD (LI-COR 925-68070, 1:15,000).

### Oxygen consumption rate (OCR) in intact cells

OCR was accessed in Seahorse XFe96 Analyzer (Agilent Technologies). VP3 96-well microplate (Agilent Technologies) were coated with poly-D-lysine (Gibco^TM^) following the manufacturer’s protocol. Three to four independently growing flasks of each genotype (WT, IF1-KO, IF1-RSC, and IF1-KO UCP4) with 60-80% confluency cells were trypsinized and cell density estimated in Bio-Rad TC20 automated cell counter. 2×10^4^ cells were plated per well and incubated at 37 °C in a 5% CO_2_ humidified incubator for 18 h. Cartridge was hydrated according to the manufacturer’s instruction. Cells were washed twice by dilution with XF Assay medium, supplemented with D-glucose 10 mM, pyruvate 1 mM, and glutamine 2 mM. Cells were incubated in a CO_2_-free BOD type incubator at 37 °C for 1 h. MitoStress protocol was used to access mitochondrial function. Final concentration of inhibitors was: oligomycin 1 μM (Port A), FCCP 1 μM (Port B), and rotenone 1 μM plus antimycin A 1 μM (Port C) (all Sigma-Aldrich®). At the end of the assay, medium was carefully removed, and cells were washed with ice-cold PBS. PBS was removed and cells were lysed with 20 μL of RIPA Lysis and Extraction Buffer supplemented with Halt™ Protease and Phosphatase Inhibitor Cocktail (Thermo Scientific^TM^). Protein content was estimated using Pierce™ BCA Protein Assay Kits (ThermoFisher™). OCR values was corrected by protein content of each given well. The average of 6-8 replicates from the original independently growing flask was considered one independent sample (n=3). OCR and ECAR results were transformed in O_2_ pmol/min/mg protein and mpH/min/mg protein.

### Transmission Electron Microscopy Analysis

WT, KO and RSC cells were grown in duplicates in T175 flasks till confluency. Cells were collected by trypsinization and washed thrice in PBS. Subsequently, the cells were fixed at room temperature for 1 h in McDowell and Trump’s fixative and washed thrice in 0.1 M cacodylate buffer prior post-fixation in osmium tetroxide 1% (Electron Microscopy Sciences, Hatfield, PA). Cells were rinsed in distilled water and dehydrated in an ethanol series transitioning to acetone. The samples were then infiltrated with EPON-812 Resin (Electron Microscopy Sciences). After polymerization, blocks were trimmed and semithin sections (approximately 0.5 µm thick) were cut, mounted on glass slides, and stained with 1% toluidine blue in 1% sodium borate for quality check examination with a light microscope. After trimming, ultrathin sections (about 70-100 nm thick) were cut from selected blocks, placed onto 200 mesh copper grids and counterstained with 2% uranyl acetate and lead citrate. Digital images were captured using a Gatan Orius SC1000 camera attached to an FEI Co. Tecnai T12 transmission electron microscope operated at 80 kV. For quantification, at least 100 images for each study group were captured and EM subcellular analysis was made using Gatan Digital Micrograph software. Statistical analysis was performed using the non-parametric Mann-Whitney U test and data are presented as mean values. *P*-values <0.05 were considered statistically significant.

### Live-cell microscopic imaging of GSH:GSSG

Experiments were performed as described previously^44–46^. WT and IF1-KO HEK cells were transfected with either cytoplasm or mitochondrial matrix-targeted Grx1-roGFPE2 plasmids. Epifluorescence imaging of GSSG:GSH was carried out using a fluorescence wide field imaging system consisting of a ProEM1024 EMCCD (Princeton Instruments), fitted to Leica DMI 6000B inverted epifluorescence microscope equipped with a Sutter DG4 light source. Fluorescence excitation was achieved with a custom dichroic derived from Chroma #59022 modified to enhance short wavelength excitation and bandpass filters specific to fluorophores. Grx1-roGFP2 were imaged with dual excitation of 414/10 & 480/15nm combined with a 515/30 nm emission filter. Typical time resolution was 10s. Calibration of the probe was performed by adding 2 mM dithiothreitol (DTT) twice for minimal ratio value and after washout of DTT, 0.2 mM H_2_O_2_ for maximal ratio value. All the experiments were performed in ECM containing 0.25% BSA at 37°C.

### RNA isolation, RNA-seq and Gene Expression Analysis

RNA was extracted from 3 independent cell cultures of WT, IF1-KO, and IF1-RSC or IF1-KO-UCP4 from 3-4 independent cell cultures for RNA-seq. RNA was isolated in QIAcube using QIAshredder columns and RNAeasy Mini with DNase digestion (QIAGEN). RNA concentration was estimated in NanoDrop^TM^ 2000 and Qubit^TM^ 3 fluorometer. RNA integrity was analyzed in Agilent 2100 Bioanalyzer using Agilent RNA 6000 Nano Kit, poly-A-selected selected for RNA-seq library generation. Paired-end libraries were sequenced to 100 base pairs (bp) or single-end libraries were sequenced to 75 bp on a NovaSeq 6000 platform (Illumina, San Diego, CA). Raw fastq files were filtered to exclude reads with phred quality score <20 using the FASTX-Toolkit fastq_quality_filter command line utility (http://hannonlab.cshl.edu/fastx_toolkit/). The STAR RNA-Seq aligner was used to align quality filtered reads to the human genome (hg38) and reads with mapping quality <20 were discarded with the SAMtools view command^47,48^. Reads aligning to Gencode GRCh38 Release 32 human genes were quantified with the Subread package featureCounts utility^49^. Differential expression analysis was performed in the R statistical programming environment v4.1.2 using DeSeq2 v1.34.0 (https://www.r-project.org/, reference 50). Genes with and adjusted *p*-value < 0.05 and were classified as differentially expressed.

### DNA methylation arrays and data analyses

Genomic DNA was extracted from the independent genotypes: 1) WT, 2) IF1-KO, 3) IF1-RSC, and 4) IF1-KO UCP4. Samples were bisulfite-converted using an EZ DNA Methylation kit (Zymo Research) following the manufacturer’s protocol. Differential methylation at the CpG dinucleotide level was conducted using Human Infinium Methylation EPIC 850K v2.0 BeadChip arrays (Illumina) following the Infinium methylation protocol. Methylation analysis was performed with the R Bioconductor package ChAMP v2.22.0^51^. Raw IDAT files were loaded into R with the champ.load function and probes that failed with detection with p-value >0.01 or <3 beads in 5% of samples were removed. Multi-hit, non-CpG, Chromosome X/Y and SNP-overlapping probes were filtered prior to analysis. Beta-mixture quantile normalization (BMIQ) was applied to correct for type-II probe bias^52^. Differentially methylated probes (DMPs) were identified with the champ.DMP function which implements limma for calculation of the t-statistic and p-value for each probe^53^. For the promoter analysis, probes were extracted with the EPIC TSS1500 feature annotation. The mean β value was reported for genes with more than one probe mapping to the promoter region. Hypermethylated or hypomethylated probes were defined by a change in mean β value >0 or <0, respectively, and an adjusted *p*-value <0.05. Differentially methylated regions (DMRs) were detected by the Bumphunter algorithm as implemented in the champ.DMR function with 1000 permutations^54^. The Benjamini-Hochberg procedure was applied to adjust p-values for multiple testing correction.

### Steady State Metabolomics

WT, IF1-KO and IF1-RSC cells at 60-80% confluency from 6 (six) independently growing 100 mm dishes (60 cm^2^) were trypsinized and cell density estimated in Bio-Rad TC20 automated cell counter. 5×10^6^ cells were plates in 100 mm dishes and incubated at 37 °C and 5% CO_2_ for 6 h to allow cells to attach. Cells were then washed once with 10 mL ice-cold PBS and placed on ice. 1 mL UHPLC grade methanol:water 4:1, pre-chilled at –80 °C, was added in each dish and incubated at –80 °C for 15 min. Dishes were then placed on dry ice and cells scrapped off the dish. Lysates were transferred to pre-chilled 1.5 mL microtubes and centrifuged at 20,000 *g* a 4 °C for 10 min. 900 μL of supernatant were transferred to new 1.5 mL microtubes. Solvent was dried using Vacufuge Plus (Eppendorf©) and analytes were kept in –80 °C until Untargeted Metabolomics was performed. The samples were resuspended in acetonitrile-water (2%:98%) v/v, vortexed for 10 s, and centrifuged (Eppendorf Centrifuge 5425R) at 14,000 relative centrifugal force (rcf) for 10 min at 4°C. The soluble fraction of the extract, was transferred to 2 mL autosampler vial (12 mm x 32 mm height vial, 12 mm screw cap, and PTFE/silicone septa, Agilent) containing a microvolume insert (Agilent).

Samples were analyzed using an ultra-high performance liquid chromatograph (VanquishTM Horizon UHPLC, Thermo Scientific) coupled to a high-resolution mass spectrometer (Orbitrap FusionTM Tribrid, Thermo Scientific). EASY-Max NGTM was used as the ionization source, operated in the heated-electrospray ionization (H-ESI) configuration. The source parameters in positive ionization mode were as follows: spray voltage of +4000 V, sheath gas of 50 arbitrary units (arb), auxiliary gas of 10 arb, sweep gas of 1 arb, ion transfer tube at 325°C, vaporizer at 350 °C. The source parameters used in negative ionization mode were identical, with the exception of the spray voltage of – 3000 V. Prior to measurement, the mass spectrometer was calibrated using FlexMix (Thermo Scientific) following manufacture directions. EASY-IC™ (Thermo Scientific) was used during data collection; this is a secondary reagent ion source, yielding fluoranthene ions that are used as a lock mass to improve m/z accuracy by correcting for mass errors that result from variation in m/z measurement (e.g. scan-to-scan variation) and environmental changes (e.g. ambient temperature). Chromatographic separation was carried out on a Kinetex F5 analytical column (2.1 inner diameter, 100 mm length, 100 Å, 2.6 µm particle size, Phenomenex) with corresponding guard cartridge. The column was maintained at 30°C during separation. The Vanquish solvent pre-heater was maintained at 30°C. Gradient elution was performed after an initial period of isocratic elution using water with 0.1% acetic acid v/v (A) and acetonitrile with 0.1% acetic acid v/v (B). Separation was performed as follows: 0% B from 0 – 2.0 min, 0% to 100% B from 2.0 to 10.5 min, 100% B from 10.5 to 12.0 min, 100% to 0% B from 12.0 to 13.0 min, 0% B from 13.0 to 20.0 min. The flow rate was 500 µL min-1. A 10 µL static mixer was used. The post column flow path consisted of a ViperTM connection (0.1 mm ID, 550 mm length, Thermo Scientific) from the column to a six-port value and PEEK tubing from the six-port value to the ionization source.

LC-MS data were collected from individual samples, system blanks, and a pooled quality control (QC), which was generated by combining aliquots from every sample in the data set into one pooled sample. Samples and system blanks were each injected at 4 µL; the QC pool was injected 9 times at different volumes (3 injections each at 2 µL, 4 µL, and 6 µL). MS data were collected with an anticipated LC peak width of 8 s and a default charge of 1. MS data were acquired at 120,000 resolution from m/z 100-1000 with an RF lens of 60% and maximum injection time of 50 ms. Prior to acquiring LC-MS data for samples, liquid chromatography – tandem mass spectrometry (LC-MS/MS) data were acquired using the AcquireX (Thermo Scientific) deep scan methodology. The acquisition order for AcquireX was the following: system blank (n=1 injection) for exclusion list generation, QC (n=1 injection) for inclusion list generation, and n=7 injections of QC. MS and MS/MS data were collected with an anticipated LC peak width of 8 s and a default charge of 1. MS data were acquired at 120,000 resolution from m/z 100-1000 with an RF lens of 60% and maximum injection time of 50 ms. MS/MS data were acquired at 30,000 resolution using an isolation width of 1.5 (m/z), stepped assisted higher-energy collision induced dissociation was used with energy steps of 20, 35, and 60 normalized collision energy, and a maximum injection time of 54 ms. The inclusion list was generated and updated via AcquireX with a low and high mass tolerance of 5 part-per-million (ppm) mass error. An intensity filter was applied with an intensity threshold of 2.0 x 104. Dynamic exclusion was used with the following parameters: exclude after n = 3 times; if occurs within 15 s; exclusion duration of 6 s; a low mass tolerance of 5 ppm mass error; a high mass tolerance of 5 ppm mass error; and excluding isotopes.

Data files (.raw) were processed with Compound Discoverer 3.3.0.550 (ThermoFisher Scientific) to identify unique molecular features and, where possible, annotate them with chemical names. Features with distinct measured accurate mass, unique retention time, and MS/MS data were tabulated after removal of isotope peaks, blank contaminants, and noise artifacts from the data. Spearman’s p (evaluates a monotonic response), Pearson’s r (evaluates a linear response), and coefficient of determination (R2, evaluates fit to a linear model); for Spearman and Pearson correlations, p=0.05 was used as the statistical metric for significance. For every feature in the dataset, a value of these metrics was calculated to evaluate the signal response for that feature over the QC range. Any feature for which the value does not meet the filtering parameter, or any feature with a negative correlation (Spearman or Pearson), is filtered out of the dataset, ensuring that only features displaying positive correlations within the p=0.05 parameter are retained for further evaluation and interpretation.

### Methyl-tracing

WT and IF1-KO cells were grown for 4 (four) days in DMEM, high glucose, no glutamine, no methionine, no cystine (Gibco^TM^, cat # 21013024), supplemented with dialyzed FBS 10%, pyruvate 1 mM, uridine 50 μg/mL, glutamine 4 mM (all Gibco^TM^), cystine-HCl 0.2 mM (Thermo Scientific Chemicals, cat # J61651.09), and L-Methionine-(methyl-d3) 0.2 mM (Sigma-Aldrich, cat # 300616), 0.16% penicillin/streptomycin, hygromycin 150 μg/mL, and blasticidin 5 μg/mL. 60-80% confluent cells from 3 (three) independently growing 100 mm dishes (60 cm^2^) were trypsinized and cell density estimated in Bio-Rad TC20 automated cell counter. Metabolite extraction was performed as described above. After extraction, samples were dried down in a centrifugal vacuum concentrator and dissolved in 4/1 acetonitrile/water mixture.

Metabolite analysis was conducted on a QExactive HF-X mass spectrometer equipped with a HESI II probe. The mass spectrometer was coupled to a Vanquish binary UPLC system (Thermo Fisher Scientific, San Jose, CA). For chromatographic separation prior to mass analysis, 5 uL of the sample was injected onto a BEH Z-HILIC column (100 mm, 1.7uM particle size, 2.1 mm internal diameter, Waters). Samples were diluted 1:5 in acetonitrile for the analysis. For chromatographic separation, Mobile phase A was 15 mM ammonium bicarbonate (pH 9.0) in 90% water and 10% acetonitrile, and mobile phase B was 15mM ammonium bicarbonate (pH 9.0) in 95% acetonitrile and 5% water. Water and acetonitrile were purchased from Fisher and were Optima LC/MS grade, analytical standards were from Ammonium bicarbonate powder was purchased from Merck. The column oven was held at 40°C and autosampler at 4°C. The chromatographic gradient was carried out at a flow rate of 0.5 ml/min as follows: 0.75 min initial hold at 95% B; 0.75-3.00 min linear gradient from 95% to 30% B, 1.00 min isocratic hold at 30% B. B was brought back to 95% over 0.50 minutes, after which the column was re-equilibrated under initial conditions. Each sample was analyzed in both positive and negative modes, with the spray voltage set to 3 kV (3.5 kV for positive mode), the capillary temperature to 320 °C, and the HESI probe to 300 °C. The sheath gas flow was set to 50 units, the auxiliary gas flow was set to 10 units, and the sweep gas flow was set to 1 unit. Mass acquisition was performed in a range of m/z = 70–900, with the resolution set at 120,0000. Retention times were determined using authentic standards; in the absence of an analytical standard, a PRM experiment was conducted on candidate peaks (resolution 30,000, 1.0 Da isolation window, stepped N(CE) 30, 50, 150), and compared to publicly available spectra. Raw data were converted to mzML and processed using emzed^55^. Theoretical isotopologue m/z values for methyl-labelled metabolites were determined using the exact mass of the M0 isotopologue as well as CD3. Raw peak areas were calculated by integrating an RT window determined by m/z, chemical standards, fragmentation patterns, and isotopologue patterns.

### Sub-chronic chemical exposures

2×10^5^ cells were plated were in 6-well plates (2.2×10^4^ cells/cm^2^) and incubated for 24 h. Cells were treated with telmisartan 2 μM and annatto 10 μM for 10 days. Fresh media was added every day. After the start of the treatment, cells were sub-cultivated in the presence of the drugs. At the 10^th^ day, cells were grown in 150 mm dishes (150 cm^2^). Cells were split and plated for Seahorse analysis. The remaining pellet was collected and aliquoted for PC/PE analysis, DNA, RNA, and protein isolation.

### Estimation of cellular PE and PC content

PC and PE were estimated using phosphatidylcholine and phosphotidylethanolamine assay kit (Abcam©) according to the manufacturer’s instructions. Briefly, 2×10^6^ cells for PC and 1×10^6^ cells for PE were pelleted. PE was extracted in peroxide-carbonyl-free Triton X-100 (Sigma-Aldrich®) 5% (v/v) in deionized water. For all samples, background measurement was added for all samples to account for the intracellular levels of the products of the reaction catalyzed by PC and PE converter enzymes. Results were calculated from standard curve provided and presented as nmol of phospholipid per 1×10^6^ cells.

### Mitochondrial isolation

Mitochondria was isolated as previously described by Mori et al.^56^ with slight modifications. Briefly, WT and IF1-KO cells were grown in 2 (two) T-175 flasks until they reached 70-80% confluency. Cells were trypsinized and cell suspensions were centrifuged at 300 *g* at 4 °C for 10 min. Supernatant was removed and pellets were washed once with ice-cold MSHE buffer (mannitol 210 mM, sucrose 70 mM, HEPES-KOH 10 mM, pH 7.2, EDTA 2 mM, EGTA 1 mM, DTT 5 mM). Pellets were resuspended in 2 mL of MSHE buffer and cells were lysed using Omni Soft Tissue Tip^TM^ (Omni, Inc) on full speed for 5 s. Nuclei and membranes were pelleted by centrifugation at 700 *g* at 4 °C for 12 min. Mitochondria-enriched supernatants were transferred into new pre-chilled 1.5 mL microtubes and centrifuged at 9,000 *g* at 4 °C for 10 min. Protein content was estimated using Pierce™ BCA Protein Assay Kits (ThermoFisher™). 50 μg of mitochondrial fractions aliquoted in 0.6 mL microtubes, flash-frozen in liquid nitrogen and stored at –80 °C.

### Statistical Analysis

Where not specified, statistical analysis was performed using GraphPad Prism version 9.0. WT and IF1-KO were compared using Student’s t test. When comparison involved more than one group, one-way ANOVA with Dunnett’s or Tukey’s post test were performed.

## References

1. Di Lisa, F., Blank, P.S., Collona, R., Gambassi, G., Silverman, H.S., Stern, M.S., and Hansford, R.G. (1995). Mitochondrial membrane potential in single living adult rat cardiac myocytes exposed to anoxia or metabolic inhibition. The Journal of Physiology 486, 1–13. 10.1113/jphysiol.1995.sp020786.

2. Kulawiak, B., Hopker, J., Gebert, M., Guiard, B., Wiedemann, N., and Gebert, N. (2013). The mitochondrial protein import machinery has multiple connections to the respiratory chain. Biochim Biophys Acta 1827, 612–626. 10.1016/j.bbabio.2012.12.004.

3. Neupert, W., and Herrmann, J.M. (2007). Translocation of proteins into mitochondria. Annu Rev Biochem 76, 723–749. 10.1146/annurev.biochem.76.052705.163409.

4. Gunter, T.E., Yule, D.I., Gunter, K.K., Eliseev, R.A., and Salter, J.D. (2004). Calcium and mitochondria. FEBS Lett 567, 96–102. 10.1016/j.febslet.2004.03.071.

5. Pak, O., Sommer, N., Hoeres, T., Bakr, A., Waisbrod, S., Sydykov, A., Haag, D., Esfandiary, A., Kojonazarov, B., Veit, F., et al. (2013). Mitochondrial hyperpolarization in pulmonary vascular remodeling. Mitochondrial uncoupling protein deficiency as disease model. Am J Respir Cell Mol Biol 49, 358–367. 10.1165/rcmb.2012-0361OC.

6. Michelakis, E.D., Sutendra, G., Dromparis, P., Webster, L., Haromy, A., Niven, E., Maguire, C., Gammer, T.-L., Mackey, J.R., Fulton, D., et al. (2010). Metabolic Modulation of Glioblastoma with Dichloroacetate. Science Translational Medicine 2, 1–8. 10.1126/scitranslmed.3000677.

7. Korshunov, S.S., Skulachev, V.P., and Starkov, A.A. (1997). High protonic potential actuates a mechanism of production of reactive oxygen species in mitochondria. FEBS Lett 416, 15–18. 10.1016/s0014-5793(97)01159-9.

8. Duchen, M.R. (2000). Mitochondria and calcium: from cell signalling to cell death. J Physiol 529 *Pt* *1*, 57–68. 10.1111/j.1469-7793.2000.00057.x.

9. Panchenko, M.V., and Vinogradov, A.D. (1985). Interaction between the mitochondrial ATP synthetase and ATPase inhibitor protein. FEBS Lett 184, 226–230.

10. Pullman, M.E., and Monroy, G.C. (1963). A Naturally Occurring Inhibitor of Mitochondrial Adenosine Triphosphatase. Journal of Biological Chemistry 238, 3762–3769. 10.1016/s0021-9258(19)75338-1.

11. Chen, W.W., Birsoy, K., Mihaylova, M.M., Snitkin, H., Stasinski, I., Yucel, B., Bayraktar, E.C., Carette, J.E., Clish, C.B., Brummelkamp, T.R., et al. (2014). Inhibition of ATPIF1 ameliorates severe mitochondrial respiratory chain dysfunction in mammalian cells. Cell Rep 7, 27–34. 10.1016/j.celrep.2014.02.046.

12. Martinez-Reyes, I., Diebold, L.P., Kong, H., Schieber, M., Huang, H., Hensley, C.T., Mehta, M.M., Wang, T., Santos, J.H., Woychik, R., et al. (2016). TCA Cycle and Mitochondrial Membrane Potential Are Necessary for Diverse Biological Functions. Mol Cell 61, 199–209. 10.1016/j.molcel.2015.12.002.

13. Gorospe, C.M., Carvalho, G., Herrera Curbelo, A., Marchhart, L., Mendes, I.C., Niedzwiecka, K., and Wanrooij, P.H. (2023). Mitochondrial membrane potential acts as a retrograde signal to regulate cell cycle progression. Life Sci Alliance 6. 10.26508/lsa.202302091.

14. Ouyang, Y., Cunningham, C.N., Berg, J.A., Toshniwal, A.G., Hughes, C.E., Van Vranken, J.G., Jeong, M.-Y., Cluntun, A.A., Lam, G., Winter, J.M., et al. (2022). Phosphate Starvation Signaling Increases Mitochondrial Membrane Potential through Respiration-independent Mechanisms. BioRxiv. 10.1101/2022.10.25.513802.

15. Faccenda, D., Gorini, G., Jones, A., Thornton, C., Baracca, A., Solaini, G., and Campanella, M. (2021). The ATPase Inhibitory Factor 1 (IF(1)) regulates the expression of the mitochondrial Ca(2+) uniporter (MCU) via the AMPK/CREB pathway. Biochim Biophys Acta Mol Cell Res 1868, 118860. 10.1016/j.bbamcr.2020.118860.

16. Weissert, V., Rieger, B., Morris, S., Arroum, T., Psathaki, O.E., Zobel, T., Perkins, G., and Busch, K.B. (2021). Inhibition of the mitochondrial ATPase function by IF1 changes the spatiotemporal organization of ATP synthase. Biochim Biophys Acta Bioenerg 1862, 148322. 10.1016/j.bbabio.2020.148322.

17. Acin-Perez, R., Beninca, C., Fernandez Del Rio, L., Shu, C., Baghdasarian, S., Zanette, V., Gerle, C., Jiko, C., Khairallah, R., Khan, S., et al. (2023). Inhibition of ATP synthase reverse activity restores energy homeostasis in mitochondrial pathologies. EMBO J 42, e111699. 10.15252/embj.2022111699.

18. Muller, C.S., Bildl, W., Haupt, A., Ellenrieder, L., Becker, T., Hunte, C., Fakler, B., and Schulte, U. (2016). Cryo-slicing Blue Native-Mass Spectrometry (csBN-MS), a Novel Technology for High Resolution Complexome Profiling. Mol Cell Proteomics 15, 669–681. 10.1074/mcp.M115.054080.

19. To, T.L., Cuadros, A.M., Shah, H., Hung, W.H.W., Li, Y., Kim, S.H., Rubin, D.H.F., Boe, R.H., Rath, S., Eaton, J.K., et al. (2019). A Compendium of Genetic Modifiers of Mitochondrial Dysfunction Reveals Intra-organelle Buffering. Cell 179, 1222–1238 e1217. 10.1016/j.cell.2019.10.032.

20. Johnson, J.M., Peterlin, A.D., Balderas, E., Sustarsic, E.G., Maschek, J.A., Lang, M.J., Jara-Ramos, A., Panic, V., Morgan, J.T., Villanueva, C.J., et al. (2023). Mitochondrial phosphatidylethanolamine modulates UCP1 to promote brown adipose thermogenesis. Science Advances 9, 14. 10.1126/sciadv.ade7864.

21. Santos, J.H. (2021). Mitochondria signaling to the epigenome: A novel role for an old organelle. Free Radic Biol Med 170, 59–69. 10.1016/j.freeradbiomed.2020.11.016.

22. Lozoya, O.A., Martinez-Reyes, I., Wang, T., Grenet, D., Bushel, P., Li, J., Chandel, N., Woychik, R.P., and Santos, J.H. (2018). Mitochondrial nicotinamide adenine dinucleotide reduced (NADH) oxidation links the tricarboxylic acid (TCA) cycle with methionine metabolism and nuclear DNA methylation. PLoS Biol 16, e2005707. 10.1371/journal.pbio.2005707.

23. Lozoya, O.A., Xu, F., Grenet, D., Wang, T., Grimm, S.A., Godfrey, V., Waidyanatha, S., Woychik, R.P., and Santos, J.H. (2020). Single Nucleotide Resolution Analysis Reveals Pervasive, Long-Lasting DNA Methylation Changes by Developmental Exposure to a Mitochondrial Toxicant. Cell Rep 32, 108131. 10.1016/j.celrep.2020.108131.

24. Mao, W., Yu, X.X., Zhong, A., Li, W., Brush, J., Sherwood, S.W., Adams, S.H., and Pan, G. (1999). UCP4, a novel brain-specific mitochondrial protein that reduces membrane potential in mammalian cells. FEBS Lett 443, 326–330. 10.1016/s0014-5793(98)01713-x.

25. Kohli, R.M., and Zhang, Y. (2013). TET enzymes, TDG and the dynamics of DNA demethylation. Nature 502, 472–479. 10.1038/nature12750.

26. Cortellino, S., Xu, J., Sannai, M., Moore, R., Caretti, E., Cigliano, A., Le Coz, M., Devarajan, K., Wessels, A., Soprano, D., et al. (2011). Thymine DNA glycosylase is essential for active DNA demethylation by linked deamination-base excision repair. Cell 146, 67–79. 10.1016/j.cell.2011.06.020.

27. Tahiliani, M., Koh, K.P., Yinghua Shen, Y., Pastor, W.A., Bandukwala, H., Brudno, Y., Agarwal, S., Iyer, L.M., Liu, D.R., Aravind, L., and Rao, A. (2009). Conversion of 5-Methylcytosine to 5-Hydroxymethylcytosine in Mammalian DNA by MLL Partner TET1. Science 324, 6. 10.1126/science.1170116.

28. Xiao, M., Yang, H., Xu, W., Ma, S., Lin, H., Zhu, H., Liu, L., Liu, Y., Yang, C., Xu, Y., et al. (2012). Inhibition of alpha-KG-dependent histone and DNA demethylases by fumarate and succinate that are accumulated in mutations of FH and SDH tumor suppressors. Genes Dev 26, 1326–1338. 10.1101/gad.191056.112.

29. Xu, W., Yang, H., Liu, Y., Yang, Y., Wang, P., Kim, S.H., Ito, S., Yang, C., Wang, P., Xiao, M.T., et al. (2011). Oncometabolite 2-hydroxyglutarate is a competitive inhibitor of alpha-ketoglutarate-dependent dioxygenases. Cancer Cell 19, 17–30. 10.1016/j.ccr.2010.12.014.

30. Cyr, A.R., Hitchler, M.J., and Domann, F.E. (2013). Regulation of SOD2 in cancer by histone modifications and CpG methylation: closing the loop between redox biology and epigenetics. Antioxid Redox Signal 18, 1946–1955. 10.1089/ars.2012.4850.

31. Ye, C., Sutter, B.M., Wang, Y., Kuang, Z., and Tu, B.P. (2017). A Metabolic Function for Phospholipid and Histone Methylation. Mol Cell 66, 180–193 e188. 10.1016/j.molcel.2017.02.026.

32. Locasale, J.W. (2013). Serine, glycine and one-carbon units: cancer metabolism in full circle. Nat Rev Cancer 13, 572–583. 10.1038/nrc3557.

33. Attene-Ramos, M.S., Huang, R., Michael, S., Witt, K.L., Richard, A., Tice, R.R., Simeonov, A., Austin, C.P., and Xia, M. (2015). Profiling of the Tox21 chemical collection for mitochondrial function to identify compounds that acutely decrease mitochondrial membrane potential. Environ Health Perspect 123, 49–56. 10.1289/ehp.1408642.

34. Fang, W., Zhu, Y., Yang, S., Tong, X., and Ye, C. (2022). Reciprocal regulation of phosphatidylcholine synthesis and H3K36 methylation programs metabolic adaptation. Cell Rep 39, 110672. 10.1016/j.celrep.2022.110672.

35. Antequera, F., Tamame, M., Villanueva, J.R., and Santos, T. (1984). DNA methylation in the fungi. Journal of Biological Chemistry 259, 8033–8036. 10.1016/s0021-9258(17)39681-3.

36. Alberts, B., Heald, R., Johnson, A., Morgan, D., Raff, M., Roberts, K., and Walter, P. (2022). Molecular Biology of the Cell, 7th Edition (W. W. Norton & Company).

37. Vance, D.E. (2014). Phospholipid methylation in mammals: from biochemistry to physiological function. Biochim Biophys Acta 1838, 1477–1487. 10.1016/j.bbamem.2013.10.018.

38. Lee RG, Rudler DL, Raven SA, Peng L, Chopin A, Moh ESX, McCubbin T, Siira SJ, Fagan SV, DeBono NJ, Stentenbach M, Browne J, Rackham FF, Li J, Simpson KJ, Marcellin E, Packer NH, Reid GE, Padman BS, Rackham O, Filipovska A. (2024). Quantitative subcellular reconstruction reveals a lipid mediated inter-organelle biogenesis network. Nat Cell Biol. 2023 Dec 21. doi: 10.1038/s41556-023-01297-4.

39. Naghdi, S., P. Varnai, and G. Hajnoczky, Motifs of VDAC2 required for mitochondrial Bak import and tBid-induced apoptosis. Proc Natl Acad Sci U S A, 2015. 112(41): p. E5590–9.

40. Roy, S.S. and G. Hajnoczky, Fluorometric methods for detection of mitochondrial membrane permeabilization in apoptosis. Methods Mol Biol, 2009. 559: p. 173–90.

41. Shechter, D., Dormann, H.L., Allis, C.D., and Hake, S.B. (2007). Extraction, purification and analysis of histones. Nat Protoc 2, 1445–57. 10.1038/nprot.2007.202.

42. Jha, P., Wang, X., Auwerx, J. (2016). Analysis of Mitochondrial Respiratory Chain Supercomplexes Using Blue Native Polyacrylamide Gel Electrophoresis (BN-PAGE). Curr. Protoc. Mouse Biol. 6:1–14. 10.1002/9780470942390.mo150182

43. Brischigliaro, M., Cabrera-Orefice, A., Arnold, S., Viscomi, C., Zeviani, M., Fernández-Vizarra, F. (2023). Structural rather than catalytic role for mitochondrial respiratory chain supercomplexes. 12, RP88084. 10.7554/eLife.88084

44. Naghdi, S., et al., Mitochondrial fusion and Bid-mediated mitochondrial apoptosis are perturbed by alcohol with distinct dependence on its metabolism. Cell Death Dis, 2018. 9(10): p. 1028.

45. Booth, D.M., et al., Oxidative bursts of single mitochondria mediate retrograde signaling toward the ER. Mol Cell, 2021. 81(18): p. 3866–3876 e2.

46. Booth, D.M., et al., Fluorescence imaging detection of nanodomain redox signaling events at organellar contacts. STAR Protoc, 2022. 3(1): p. 101119.

47. Dobin, A., Davis, C.A., Schlesinger, F., Drenkow, J., Zaleski, C., Jha, S., Batut, P., Chaisson, M., and Gingeras, T.R. (2013). STAR: ultrafast universal RNA-seq aligner. Bioinformatics, 29(1), 15–21. 10.1093/bioinformatics/bts635.

48. Li, H., Handsaker, B., Wysoker, A., Fennell, T., Ruan, J., Homer, N., Marth, G., Abecasis, G., Durbin, R., and 1000 Genome Project Data Processing Subgroup. (2009). The sequence alignment/map format and SAMtools. Bioinformatics, 25(16), 2078–2079.

49. Liao, Y., Smyth, G. K., and Shi, W. (2014). featureCounts: an efficient general purpose program for assigning sequence reads to genomic features. Bioinformatics 30, 923–930, doi:10.1093/bioinformatics/btt656).

50. Love, M.I., Huber, W., and Anders, S. (2014). Moderated estimation of fold change and dispersion for RNA-seq data with DESeq2. Genome Biol 15, 550. 10.1186/s13059-014-0550-8.

51. Tian, Y., Morris, T.J., Webster, A.P., Yang, Z., Beck, S., Feber, A., Teschendorff, A.E. (2017). ChAMP: Updated methylation analysis pipeline for Illumina BeadChips. Bioinformatics 33, 3982–3984.

52. Teschendorff, A. E., Marabita, F., Lechner, M., Bartlett, T., Tegner, J., Gomez-Cabrero, D., and Beck, S. (2013). A beta-mixture quantile normalization method for correcting probe design bias in Illumina Infinium 450 k DNA methylation data. Bioinformatics, 29(2), 189–196.

53. Smyth, G. K. limma: linear models for microarray data. (2005). Bioinformatics and Computational Biology Solutions Using R and Bioconductor (eds Gentleman, R. et al.) 397–420 (Springer).

54. Jaffe, A. E., Murakami, P., Lee, H., Leek, J. T., Fallin, M. D., Feinberg, A. P., and Irizarry, R. A. (2012). Bump hunting to identify differentially methylated regions in epigenetic epidemiology studies. International journal of epidemiology, 41(1), 200–209. 10.1093/ije/dyr238.

55. Kiefer, P., Schmitt, U., Vorholt, J.A. (2013). eMZed: an open source framework in Python for rapid and interactive development of LC/MS data analysis workflows. Bioinformatics 29, 963–964.

56. Mori, M.P., Costa, R.A.P., Soltys, D.T., Freire, T.S., Rossato, F.A., Amigo, I., Kowaltowski, A.J., Vercesi, A.E., and Souza-Pinto, N.C. (2017). Lack of XPC leads to a shift between respiratory complexes I and II but sensitizes cells to mitochondrial stress. Sci Rep. 7, 155. 10.1038/s41598-017-00130-x.

