## Supplemental figures for "Mitochondrial membrane potential regulates nuclear DNA methylation and gene expression through phospholipid remodeling"

Figure S1

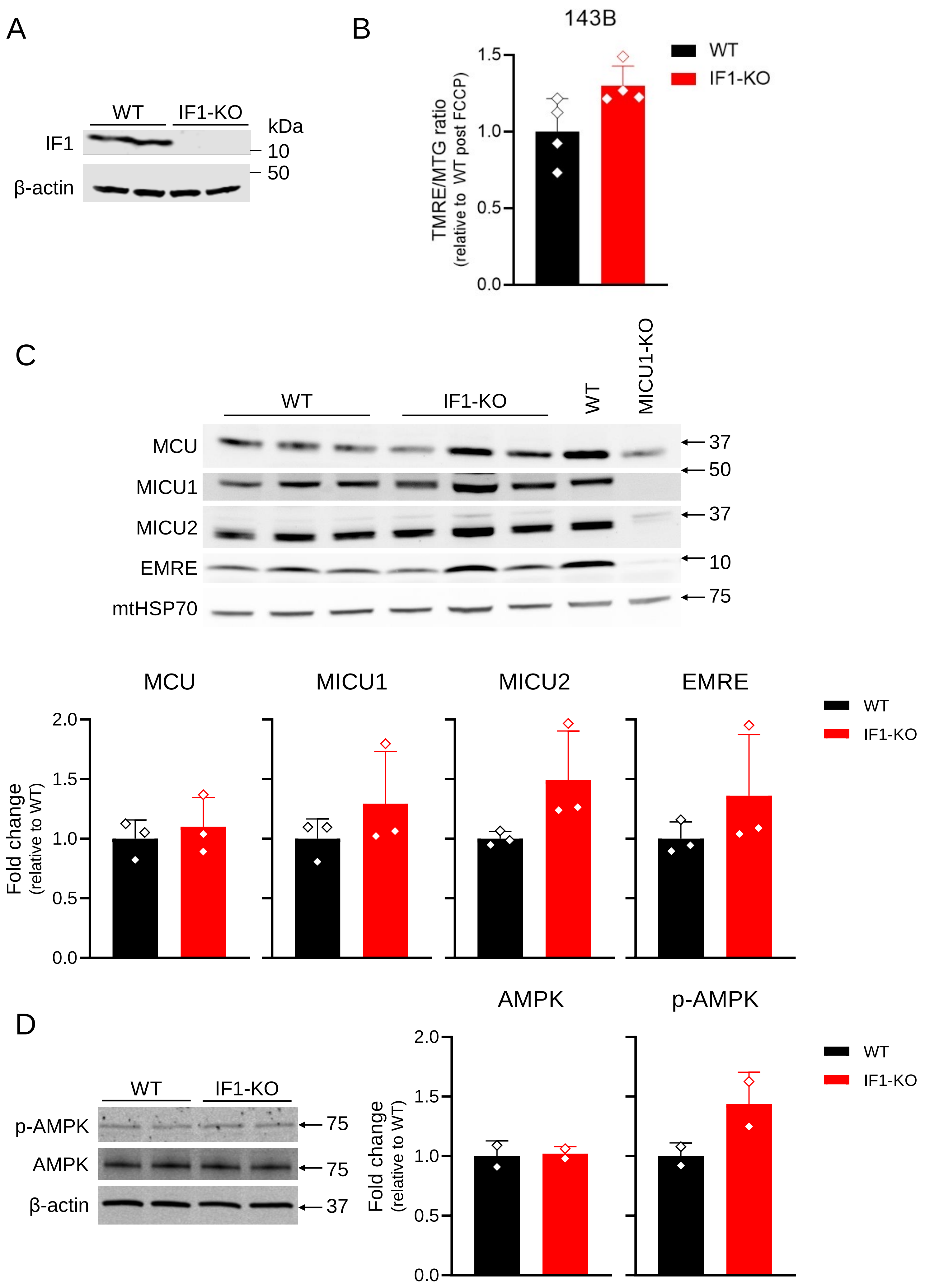

A
WT
IF1-KO
kDa
IF1
10
50
β-actin
B
143B
WT
IF1-KO
TMRE/MTG ratio
(relative to WT post FCCP)
1.5
1.0
0.5
0.0
C
WT
IF1-KO
WT
MICU1-KO
MCU
MICU1
MICU2
EMRE
mtHSP70
37
50
37
10
75
MCU
MICU1
MICU2
EMRE
WT
IF1-KO
Fold change
(relative to WT)
2.0
1.5
1.0
0.5
0.0
D
WT
IF1-KO
p-AMPK
AMPK
β-actin
75
75
37
AMPK
p-AMPK
WT
IF1-KO
Fold change
(relative to WT)
2.0
1.5
1.0
0.5
0.0

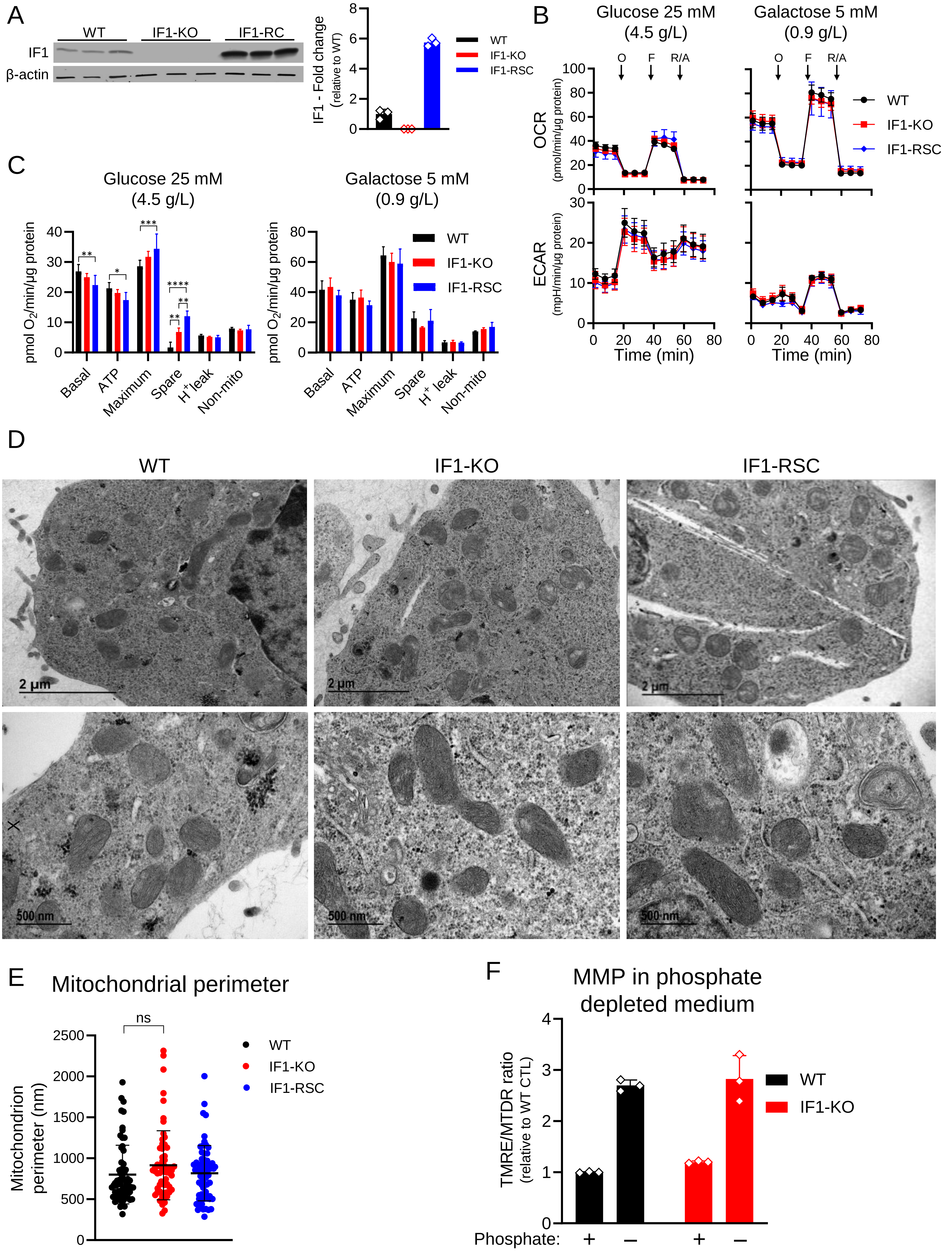

A
WT
IF1-KO
IF1-RC
IF1
β-actin
IF1 - Fold change
(relative to WT)
WT
IF1-KO
IF1-RSC
B
Glucose 25 mM
(4.5 g/L)
Galactose 5 mM
(0.9 g/L)
O
F
R/A
OCR
(pmol/min/μg protein)
ECAR
(mpH/min/μg protein)
Time (min)
WT
IF1-KO
IF1-RSC
C
Glucose 25 mM
(4.5 g/L)
Galactose 5 mM
(0.9 g/L)
pmol O2/min/μg protein
Basal
ATP
Maximum
Spare
H+ leak
Non-mito
WT
IF1-KO
IF1-RSC
D
WT
IF1-KO
IF1-RSC
18500X
49000 X
2 μm
500 nm
E
Mitochondrial perimeter
ns
Mitochondrion
perimeter (nm)
WT
IF1-KO
IF1-RSC
F
MMP in phosphate
depleted medium
TMRE/MTDR ratio
(relative to WT CTL)
WT
IF1-KO
Phosphate:
+
–
+
–

Figure S3

A

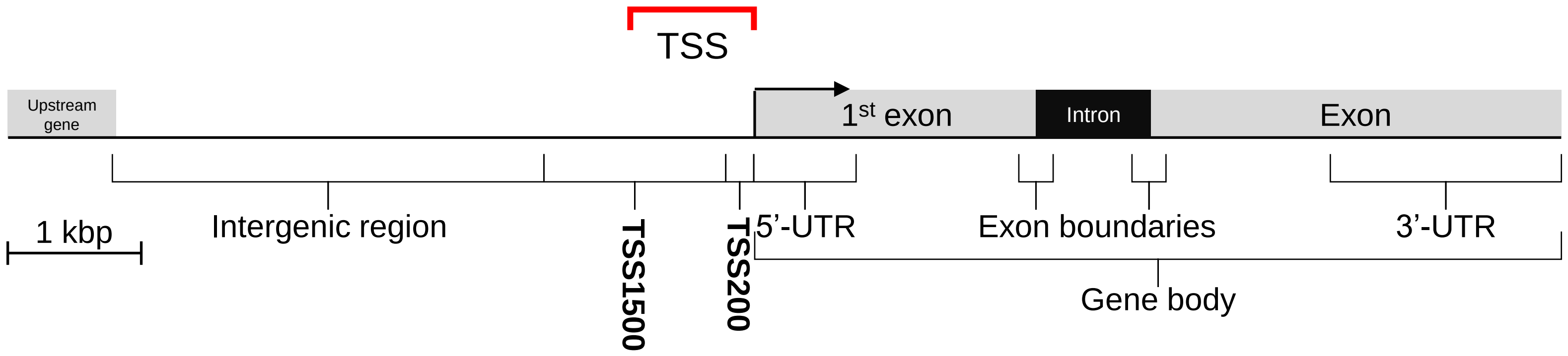


B

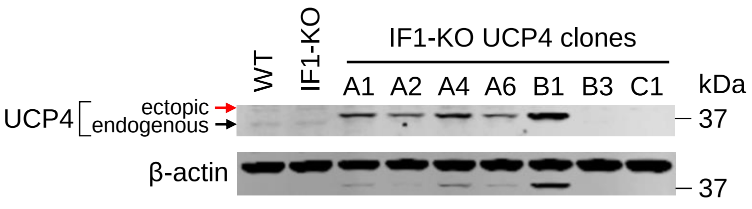


C

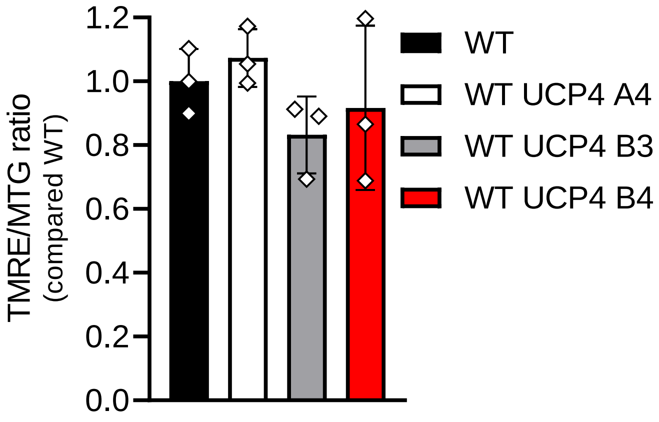


D

DNA methylation changes in RSC and UCP4 cells vs KO

| | Rescued | | | UCP4 | | |
|---|---|---|---|---|---|---|
| Methylation | Hyper | Hypo | All | Hyper | Hypo | All |
| Full | 1130 | 857 | 1987 | 1456 | 633 | 2089 |
| Partial | 2731 | 667 | 3398 | 1729 | 990 | 2719 |
| No | 4391 | 1816 | 6207 | 5161 | 1860 | 7021 |
| Somewhat | 3861 | 1524 | 5385 | 3185 | 1623 | 4808 |
| Not full | 7122 | 2483 | 9605 | 6890 | 2850 | 9740 |

### Figure S4

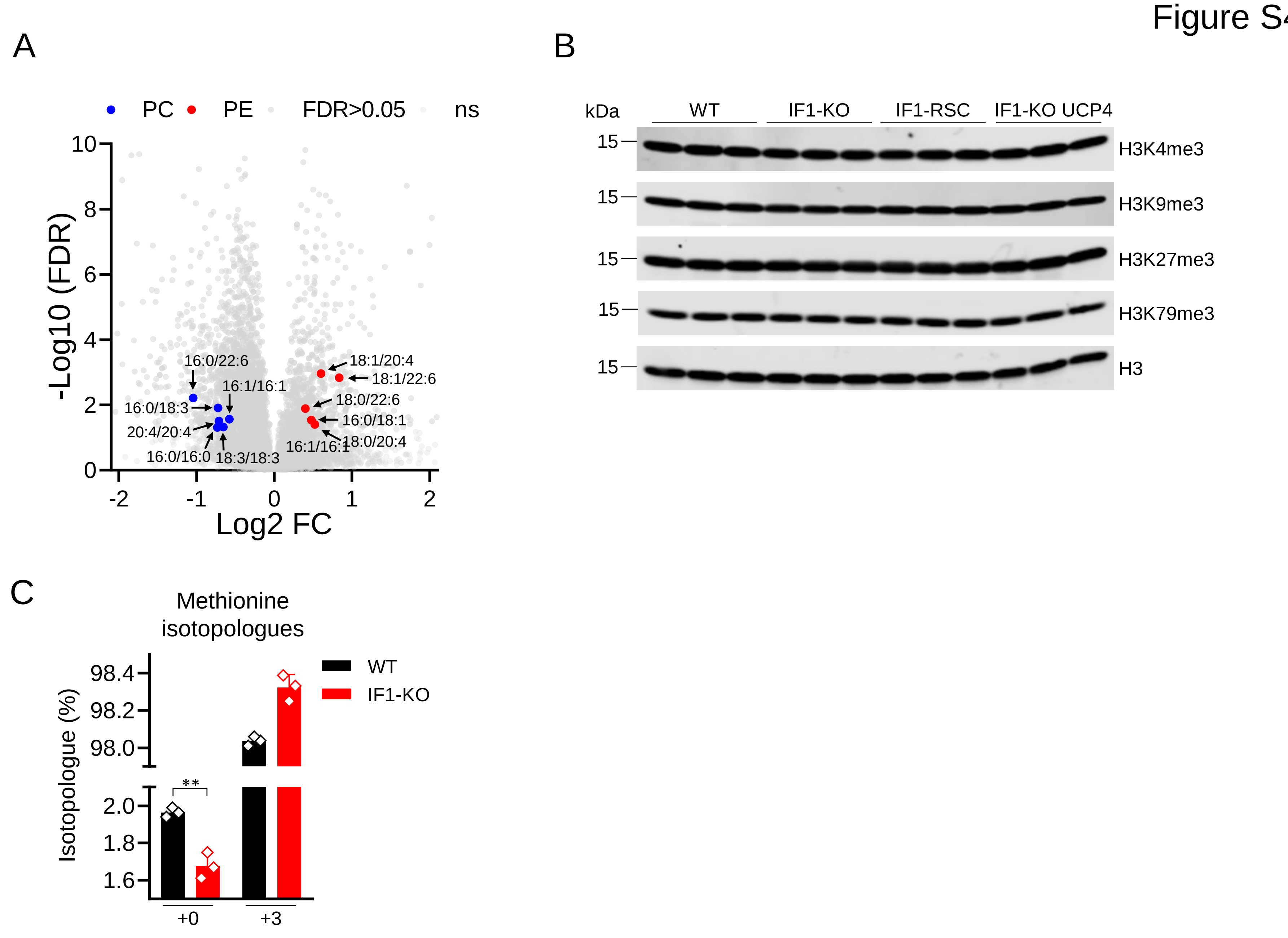

A
PC
PE
FDR>0.05
ns
-Log10 (FDR)
Log2 FC
16:0/22:6
16:1/16:1
16:0/18:3
20:4/20:4
16:0/16:0
18:3/18:3
18:1/20:4
18:1/22:6
18:0/22:6
16:0/18:1
16:1/16:1
18:0/20:4
B
kDa
WT
IF1-KO
IF1-RSC
IF1-KO UCP4
15
H3K4me3
H3K9me3
H3K27me3
H3K79me3
H3
C
Methionine isotopologues
Isotopologue (%)
WT
IF1-KO
**
+0
+3

Figure S5

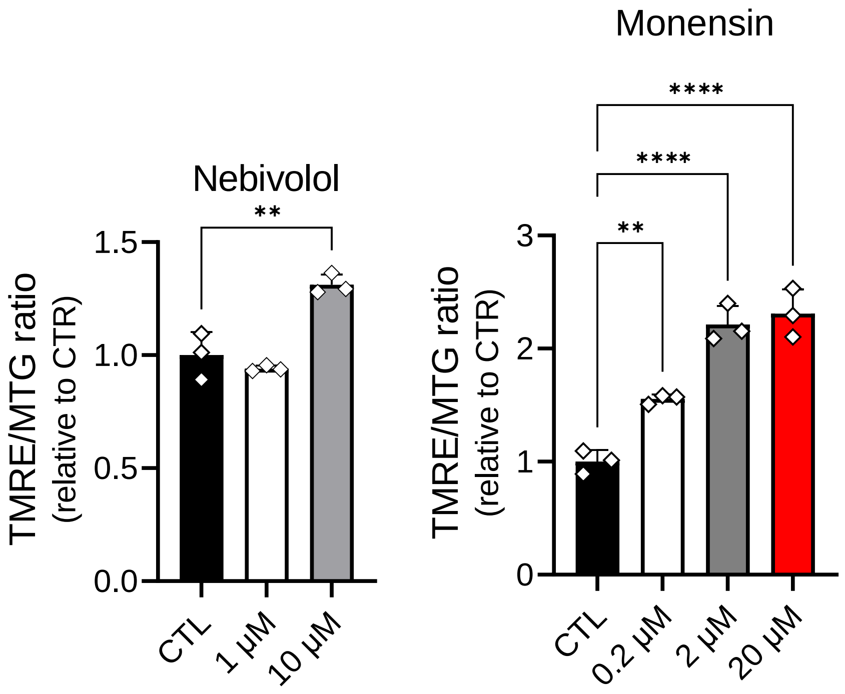

Nebivolol
Monensin
TMRE/MTG ratio
(relative to CTR)
1.5
1.0
0.5
0.0
3
2
1
0
**
****
****
**
CTL
1 µM
10 µM
CTL
0.2 µM
2 µM
20 µM
